# Bacterial lipoxygenases are associated with host-microbe interactions and may provide cross-kingdom host jumps

**DOI:** 10.1101/2022.06.21.497025

**Authors:** Georgy Kurakin

**Affiliations:** Pirogov Russian National Research Medical University, 117997, Moscow, Russia

**Keywords:** **<u>Keywords</u>**: lipoxygenases, oxylipins, cross-kingdom pathogens, cross-kingdom host jumps, emerging pathogens, cystic fibrosis

## Abstract

In this bioinformatic research, we studied the association of bacterial lipoxygenases (LOXs) with pathogenic and symbiotic traits by text networks analysis, phylogenetic analysis, and statistical analysis of molecular structure. We found that bacterial lipoxygenases are associated with a broad host range — from coral to plants and humans. In humans, bacterial LOXs are associated with opportunistic and nosocomial infections as well as with affecting specific patient populations like cystic fibrosis patients. Moreover, bacterial LOXs are associated with plant-human (or human-plant) host jumps in emerging pathogens. We also inferred a possible mechanism of such host jumps working *via* a host’s oxylipin signalling “spoofing”.

**Graphical abstract:** 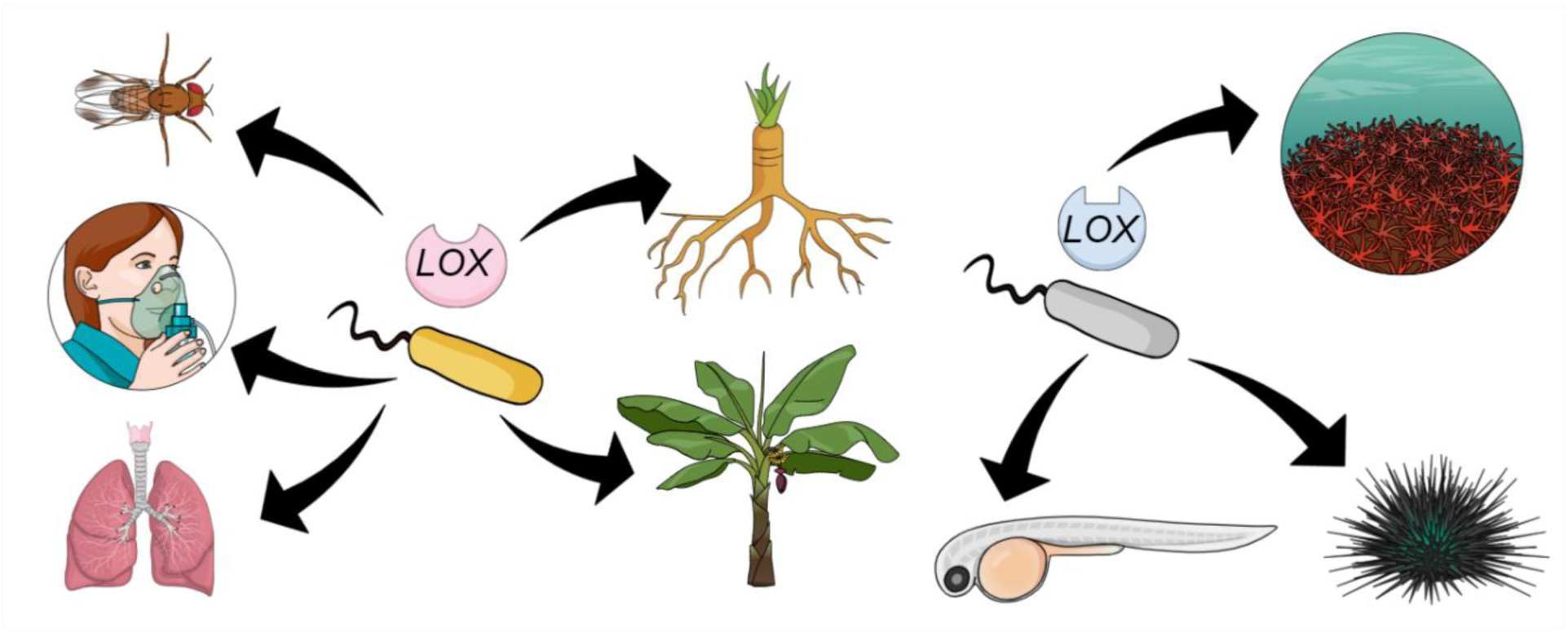

## Background

Recently, the problem of host switch (also host shift) in pathogens has gained new relevance due to the growing threat posed by emerging pathogens [1]. The mechanism of host switching by the viruses, responsible for the current (coronaviruses) or possible (henipaviruses, filoviruses) pandemics, has been particularly actively studied. Host shifts typically involve two or more relatively closely related taxa with similar morphology and physiology. Coronaviruses and filoviruses pass to humans from bats, most often via intermediate hosts (palm civets, camels, primates) [2], while new human-threatening influenza viruses often originate in waterfowl [3]. In contrast, there are virtually no known examples of viruses capable of transferring from humans to plants and *vice versa*. The only possible candidate for such a virus is the hepatitis D virus [4], which is similar to plant viroids, but direct relatedness between them is controversial.

However, cross-kingdom host jump — a host switch in which the new host belongs to a different kingdom from the previous one — is common among bacteria. Most of the known cross-kingdom jumps occur exactly between plants and humans, and such bacteria have a specific clinical profile [5]. As a rule, they cause opportunistic infections (septicemia, pneumonia, surgical infections) without affecting healthy immunocompetent individuals. Less frequently, plant-animal host jumps have been recorded [6].

The mechanisms of such host jumps are intriguing. Such distant hosts have different molecular structures determining susceptibility to infection, and the biochemical basis of pathogen compatibility to different hosts is yet unclear although extensively studied. Proteases, phospholipases, toxins, and secondary metabolism regulators have been identified as virulence factors common to plant-and animal-targeted virulence. Quorum sensing systems are also discussed in this regard [5, 7].

In our recent paper [8], we had suggested the role of lipoxygenases and oxylipin signalling in cross-kingdom pathogenicity. Lipoxygenases are very conserved enzymes involved in the synthesis of oxylipins — signalling compounds regulating stress and immune responses in a wide range of multicellular eukaryotes, such as animals (including humans), plants, and different groups of algae. The main goal of our work was to trace evolutionary origins of oxylipin signalling, using lipoxygenases as a bioinformatic proxy for oxylipin biosynthetic ability. We have found that in bacteria, lipoxygenases were primarily associated with multicellularity (at least at one stage of the life cycle) which reflects the possible role of oxylipins as ancient cell-to-cell signalling compounds.

However, some bacteria from our dataset did not possess any (albeit primitive) multicellularity. For some of these bacteria, ecophysiological data was not available. Regarding the fraction with described ecophysiology, it could be divided into three groups: (1) human-or vertebrate-associated bacteria, (2) plant-associated bacteria, (3) and bacteria associated with marine organisms. Notably, the first two groups were significantly overlapped: at least two bacteria in our dataset (*Pseudomonas aeruginosa* and *Burkholderia gladioli*) were cross-kingdom pathogens capable of infecting both plants and humans. Continued updating of our dataset resulted in adding one more species (*Pantoea ananas*) to the list of lipoxygenase-positive cross-kingdom pathogens [9]. These facts lead to an assumption that lipoxygenases may facilitate cross-kingdom host jumps and serve as versatile virulence/symbiosis factors.

In our previous paper [8], we have noticed some additional traits common for lipoxygenase-positive bacteria. They were opportunistic pathogens (these bacteria are prone to affect immunocompromised people, especially patients with cystic fibrosis). The most concerning common trait we noticed is multiple drug resistance [9].

However, these traits were noticed during the non-systematic review of literature data on bacteria in which lipoxygenases were detected. The list of these characteristic traits was not reinforced by thorough statistical analysis. The main focus of the previous papers was a link between LOXs and multicellularity, and conclusions on a link between LOXs and host-microbe interactions were preliminary.

In the current article, we continue the same research project and elaborate this association in more details. The aim of this bioinformatic research was to find additional evidence that lipoxygenases and oxylipins could contribute to host colonization and invasion, as well as to cross-kingdom host jumps.

## Materials and Methods

### Collecting and updating the dataset

We started with the same dataset of bacterial lipoxygenases as used in our previous research [8]. However, we are updating it in an ongoing manner because LOX-like sequences are constantly added to protein databases, and some sequences are revised and deleted.

The method of updating the dataset is the same as for collecting the initial dataset (described in our previous paper [8]). The difference was that we used pathogen and symbiont LOX sequences form the “old” dataset as queries and downloaded all hits that belong to pathogens and symbionts (they were typically sorted at the top of the hit list) along with some cyanobacterial and myxobacterial LOXs (according to our previous research, they are outgroups for all pathogen and symbiont LOXs). The methods for checking LOX-like hits were the same as described in our previous paper [8].

The resulting list of LOX-carrying bacteria was used for the estimation of a statistically reinforced ecological profile of bacterial LOXs and LOX-carrying bacteria. The criteria of inclusion into this list were:

1. the presence of a species in the updated LOX dataset;
2. availability of any ecological data for in the literature;
3. pathogenic/symbiotic properties or other association with any host.

### Ecological profile generation by network text analysis

We used a combination of systematic literature review and network text analysis to estimate the overall ecological profile of LOX-carrying pathogenic and symbiotic bacteria.

The species names from the list of LOX-carrying bacteria were used as the queries for PubMed database search (https://pubmed.ncbi.nlm.nih.gov/). The first ten results for each query were used for further text analysis. We used article abstracts, titles, and keywords for manual extraction of terms characterizing the ecology of the respective species (referred to below as “terms”). Totally, this research used an array of 130 papers for meta-analysis [10–138].

These terms were manually normalized to the root form and, in some cases, to the most frequent form (e.g., “emergent” to “emerging”, “growth promotion” to “growth-promoting”). An important exclusion (that could potentially bias the statistics of the term) was any terms signing antimicrobial resistance: they all were normalized as “AMR”, and any terms signing multidrug resistance were normalized as “MDR”.

For each term, we counted the number of species in which it occurs to estimate the abundance of each term in the collective ecological profile of the LOX-carrying bacteria. We excluded the terms occurring in only one species as the least representative. Then we explored associations between the terms by building a graph where each node (entity) represents a term, and each edge represents a co-occurrence of two terms (represented by its nodes) in the same species. The weight of each edge represents the number of species where the respective terms co-occur. We classified all terms to 6 groups:

1. “vertebrate-related” or “human-related” terms associated with affecting humans and pathogenicity to humans and/or vertebrates (i.e., “human”, “lung”, “abscess”, cystic fibrosis” etc.). Interactions with planktonic larval forms of fishes were excluded from this group;
2. “plant-related” terms associated with plant pathogenicity or plant symbiosis (i.e., “plant”, “rhizosphere”, “root” etc.);
3. “insect-related” group included only one term (“insect”);
4. “marine-related” terms reflecting association with marine organisms;
5. “public health threat” group (i.e. “emerging”, “AMR”, “MDR”, “threat” etc.);
6. other terms not fitting in any of the above groups.

This analysis was performed with the aid of Microsoft Excel 2019, Python 3, Gephi 0.9 [139], and Cosmograph (https://cosmograph.app/) software.

Similar ecological classification (“vertebrate-related”, “plant-related” and “marine-related”) was applied to the bacterial species for further analysis. There was a difficulty classifying bacteria which were associated with fishes. On the one hand, fishes are vertebrates, and fish-related bacteria could be considered as vertebrate-related. On the other hand, they belong ecologically to aquatic environment, and bacteria associated with marine fishes could be considered as marine-related. Here, we used additional taxonomic data for additional clarification. If the nearest related species of a bacterium affected other vertebrates (including terrestrial ones) and/or humans, this species was considered as “human/vertebrate-related”. If they affected marine invertebrates, this species was considered as a representative of “marine-related” group.

### Phylogenetics and conservation analysis

Phylogenetic analysis was performed with the same software and with the same protocols as described in our previous paper: MAFFT online v. 7 [140], MEGA X [141], iTOL [142] for phylogenetic trees and SplitsTree for phylogenetic networks [143]. Amino acid conservation analysis was performed with ConSurf server [144] in ConSeq mode [145].

### Binding site statistical analysis

For the statistical analysis of binding site structures, we used amino acid residue volumes provided by Perkins (1986) [146]. Statistical analysis was performed with MS Excel 2019 and PAST [147].

The inclusion criteria for the statistical analysis were:

1. a LOX is represented on any phylogenetic tree;
2. (preferable) the bacteria, possessing this LOX, is described in terms of their ecophysiology.

## Results

### Collecting and updating the dataset

The full updated list of LOX-carrying bacteria is provided in **Table 1** with the color-coded ecological functions. The list of bacterial species for the network text analysis along with corresponding accession IDs of lipoxygenases and PubMed IDs (PMIDs) of articles included in the analysis is provided in **Table S1**.

**Table 1.**
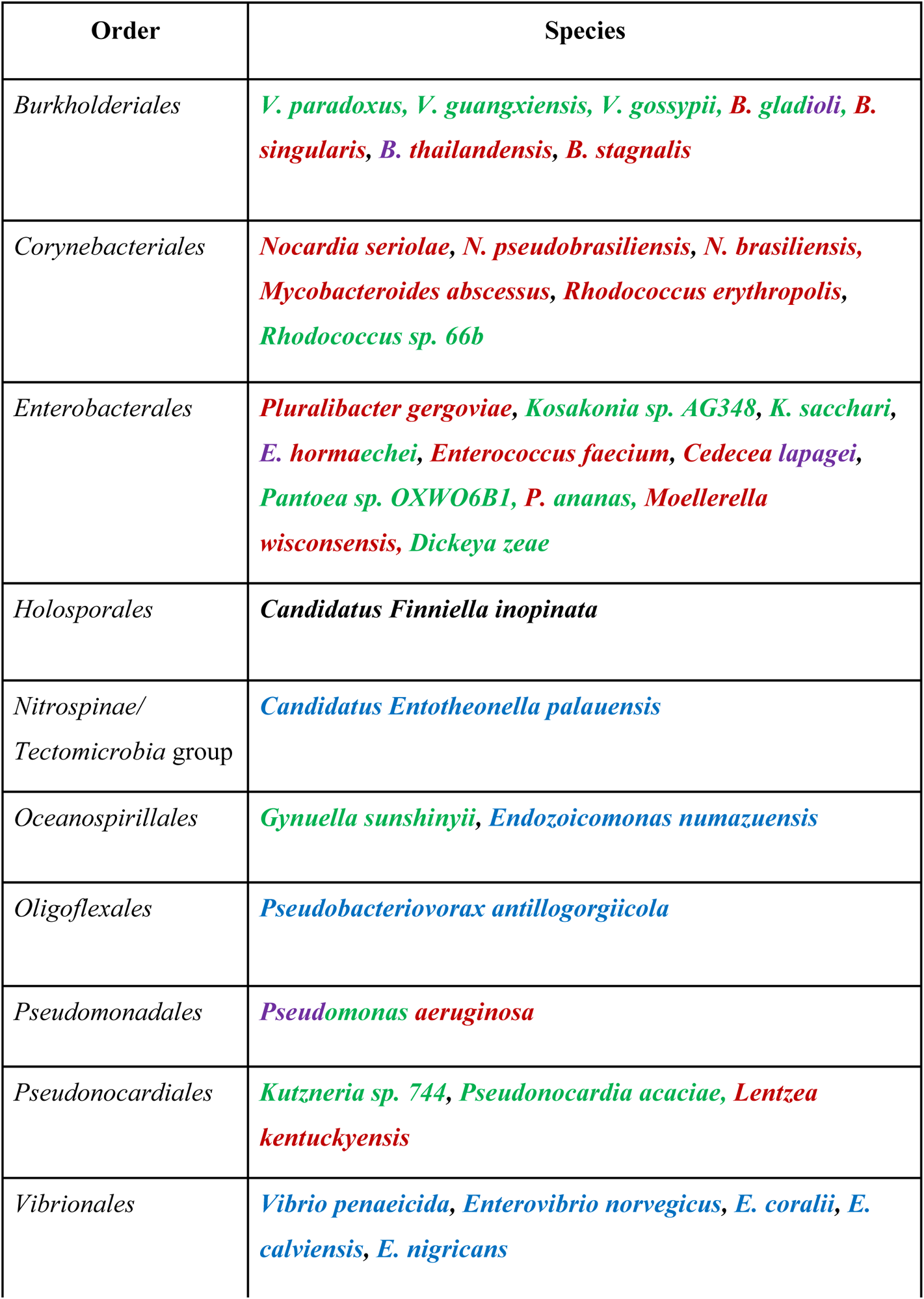
Full list of analyzed LOX-carrying species with color-coded ecological functions. Human/vertebrate-associated bacteria are depicted in **red**, plants-associated bacteria are depicted in **green**, insect-associated bacteria are depicted in **violet**, marine-associated bacteria are depicted in **blue**. If a bacterium has several host types, the corresponding colours are all represented in its names.

### The ecological profile reveals intriguing traits of LOX-carrying bacteria

The most common terms, defined as the terms with the largest number of species in which they occur (summarized in **Figure 1**, the full list provided in **Table S2**), provide some insights into the most characteristic ecological traits of LOX-carrying pathogens and symbionts. The most concerning fact is that the “public health threat” group is extremely prevalent among these bacteria. The leader of prevalence was the “AMR” term. The analysis of the most prevalent terms of the “human-related” and “plant-related” groups shed some light to some ecological details of host-microbe interaction in LOX-carrying bacteria. For instance, plant-associated LOX carriers usually colonize roots or are endophytic. LOX carriers pathogenic for humans are prone to affect lungs and cause nosocomial/opportunistic infections.

**Figure 1.**
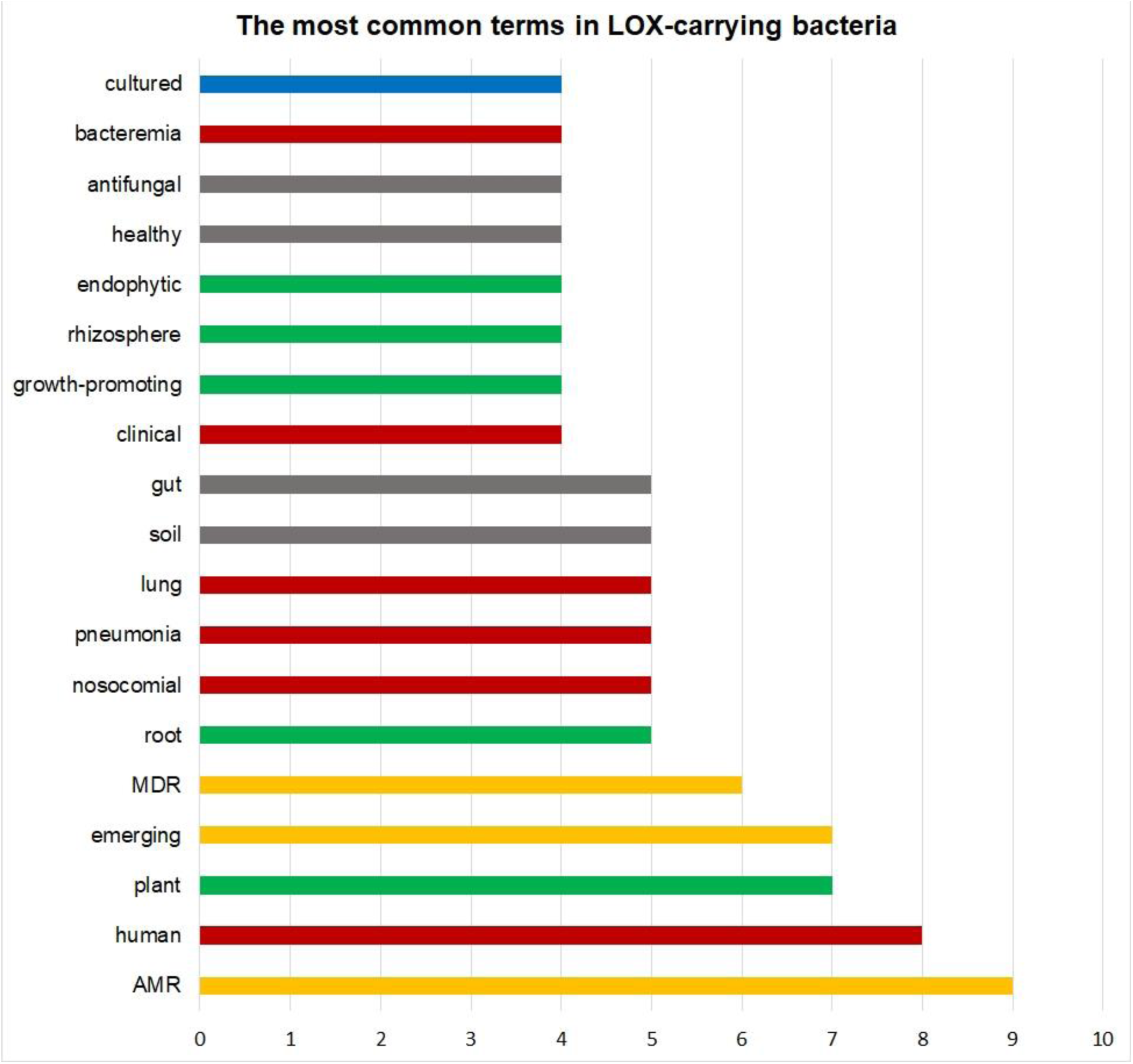
The diagram depicts the occurrence top terms (i.e., the number of species in which it occurred). This occurrence ranged from 2 to 10. Here, the most common terms are summarized, whose occurrence is not less than 4. The “human-related” and “vertebrate-related” terms are depicted in **red**, the “plant-related” terms are depicted in **green**, “public health threat” group of terms is depicted in **yellow**, and the “marine-related” terms are depicted in **blue**. *Created with MS Excel 2019*

The network analysis results (**Figure 2a, b**) show that the entire network can be divided in some clusters of terms:

1. a “plant-related” cluster comprising “plant-related” terms;
2. a “human and public health threat” cluster comprising both “human related” and “public health threat” term groups;
3. a “marine-related” cluster comprising “marine-related” terms;

The only “insect-related” term was tightly clustered with “human-related” terms suggesting the common mechanisms of affecting insects and humans.

**Figure 2a.**
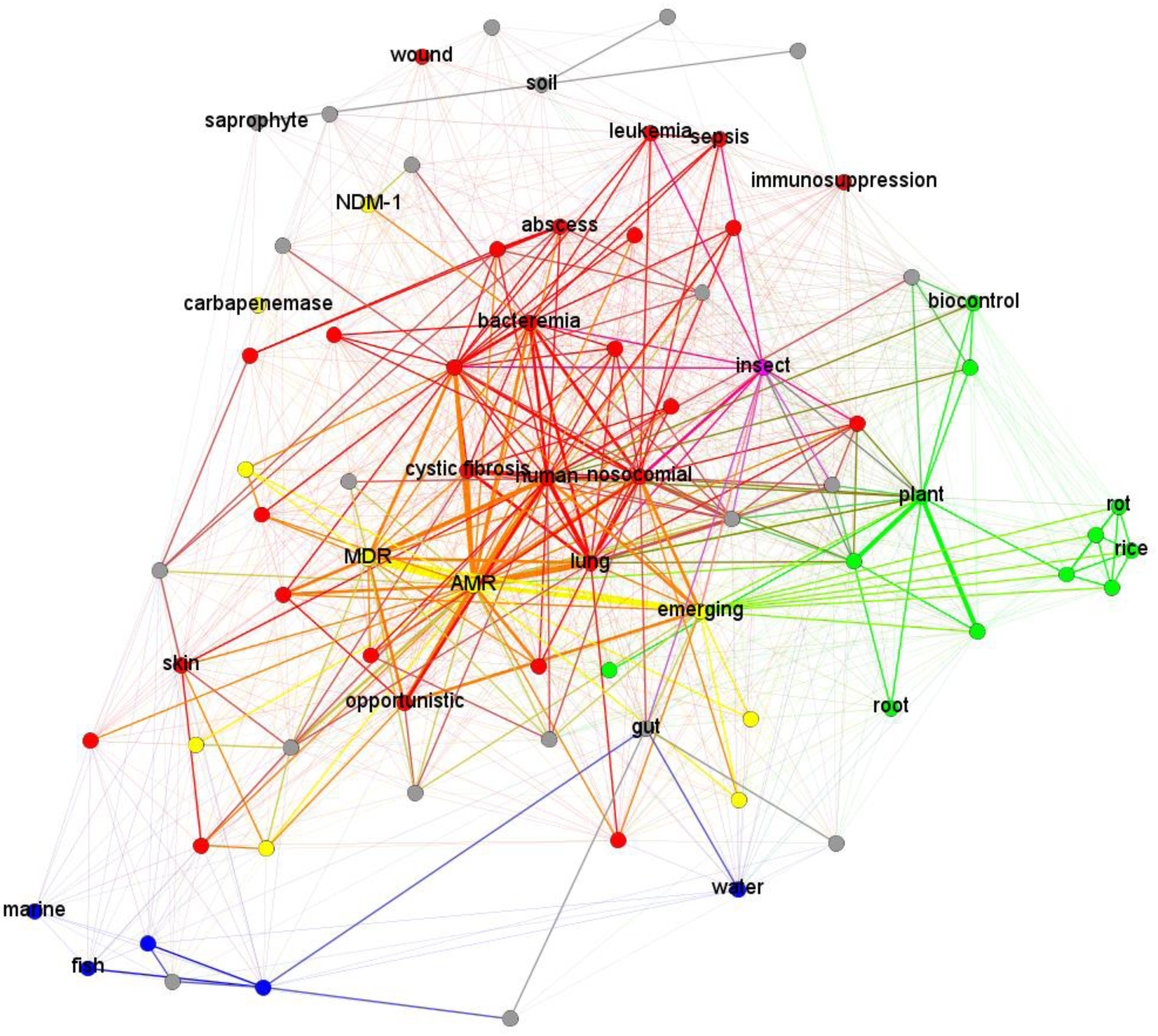
The full network of all terms in our dataset. “insect-related” terms are **magenta**, and the “marine-related” terms are **blue**. *Created with Gephi 0.9* related” group terms”. The nodes depict terms, the edges depict their occurrence in the same bacterial species. The “human-related” and “vertebrate-related” terms are depicted in **red**, the “plant-related” terms are depicted in **green**, the “public health threat” term group is depicted in **yellow**, the “insect-related” terms are **magenta**, and the “marine-related” terms are **blue**. *Created with Gephi 0.9*

**Figure 2b.**
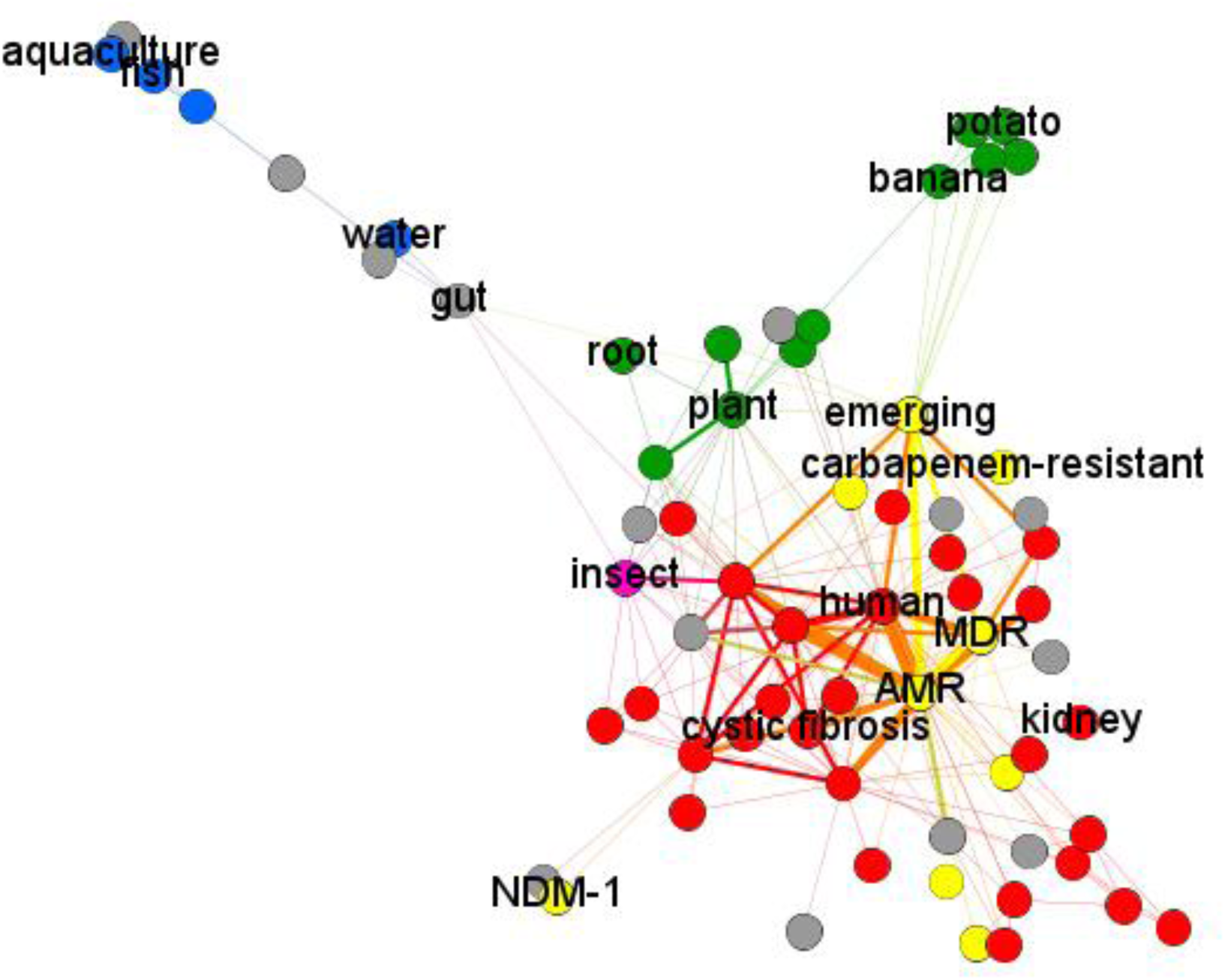
The “backbone” of our term network depicting only edges with the weight not less than 2 and the corresponding nodes only (i.e., only the terms occurring in the same LOX-carrying species not less than two times). This graph better depicts the network-forming connections and hubs of the network. Notably, the yellow nodes (the “public health threat” group terms) are the network-forming hubs providing connectivity to the entire networks, as well as the “human-related” group terms”. The nodes depict terms, the edges depict their occurrence in the same bacterial species. The “human-related” and “vertebrate-related” terms are depicted in **red**, the “plant-related” terms are depicted in **green**, the “public health threat” term group is depicted in **yellow**, the “insect-related” terms are **magenta**, and the “marine-related” terms are **blue**. *Created with Gephi 0.9*

The “plant-related” cluster had many connections to the “human and public health threat” cluster presumably *via* the network-forming hubs. These hubs represented the same terms from the “human-related” and the “public health threat” groups that were included in **Figure 1**. Thus, the “AMR” term had direct connections to 82% of all nodes including more than 50% of “plant-related” terms. The “nosocomial” term had direct connections to 68% of all nodes including the same proportion of “plant-related” terms and is tightly clustered with the “emerging” term. This fact corresponds to the most common term prevalence and confirms the fact that the human infections caused by LOX-carrying bacteria have predominantly an “emerging” status. Moreover, the term “emerging” is a hub connected to 73% of all nodes including almost all “human-related” and “plant-related” clusters (but almost not connected to the “marine-related” cluster) and reflecting the fact that pathogenic LOX carriers are predominantly characterized in the literature as emerging pathogens.

When analyzing the more specific terms, it appeared that the “cystic fibrosis” term had connections to 55% of all nodes, including 42% of “plant-related” terms such as “endophytic” and “leaves”. It is a very interesting association: the endophytic dwelling capacity in plants is associated with affecting cystic fibrosis patients in LOX-carrying bacteria. The organ-denominating term “lung” was also a hub having connections to 70% of all plant organ terms such as “leaves” and “root”. So, the same bacterial LOXs are probably associated with affecting lungs in humans and affecting roots in plants.

The results of this analysis also show that the plant pathogenesis terms (such as “rot”) and plant symbiosis terms (such as “growth-promoting”) lie within the same “plant-related” cluster. This shows that bacterial LOXs are associated both with plant pathogenicity and plant symbionts. In contrast, we do not see such an association in the “human” cluster where all terms are pathogenesis-related.

We may also conclude that this network analysis strongly supports our previous hypothesis that cross-kingdom host jump ability is a common trait of the LOX-carrying bacteria. They are mainly represented by plant-human host jumps. Conversely, “marine-related” cluster occupied the peripheral position and had a significantly lower number of connections to other clusters indicating that plant-marine or human marine host jumps are less common. It is particularly evident when analyzing only the graph “backbone” consisting of edges with weight more than or equal to 2 (**Fig. 2b**). The only “backbone” hub connecting the “marine-related” cluster with all other clusters is “gut”. This fact represents that both in marine animals and in terrestrial animals, LOX-carrying bacteria tend to dwell in gut (intestines). It is a common trait of many bacteria, and its relevance to lipoxygenases is dubious. So, “marine-related” ecological functions of LOX-carrying bacteria seem to be largely isolated from other ecological functions — such as colonizing plants, insects, and vertebrates.

The full list of term co-occurrences is provided in **Table S3**.

### Phylogenetic analysis results correspond with the collective ecological profile of LOX-carrying bacteria

The most important finding was that the results of bacterial LOX phylogenetic analysis corresponded not with the phylogeny of the bacterial species themselves rather with their ecological profile described above.

All LOXs analyzed in the current study can be divided into 4 clusters. Two clusters, “Cluster 1” and “Cluster 2”, (**Figure 3**, **Figure S1**) comprised the LOXs from phylogenetically distinct species that are: (1) human/vertebrate pathogens; (2) plant pathogens; (3) both — are able to affect both plants and humans. They reflected two independent series of the ancient horizontal transfers of the LOX genes between plant symbionts, versatile (plant/human) pathogens (including *Pseudomonas aeruginosa*), and human/vertebrate pathogens. Thus, these clusters share the common ecological function pattern with the networks in the previous section where the terms for plant pathogenicity, plant symbiosis, and vertebrate pathogenicity are tightly connected.

**Figure 3.**
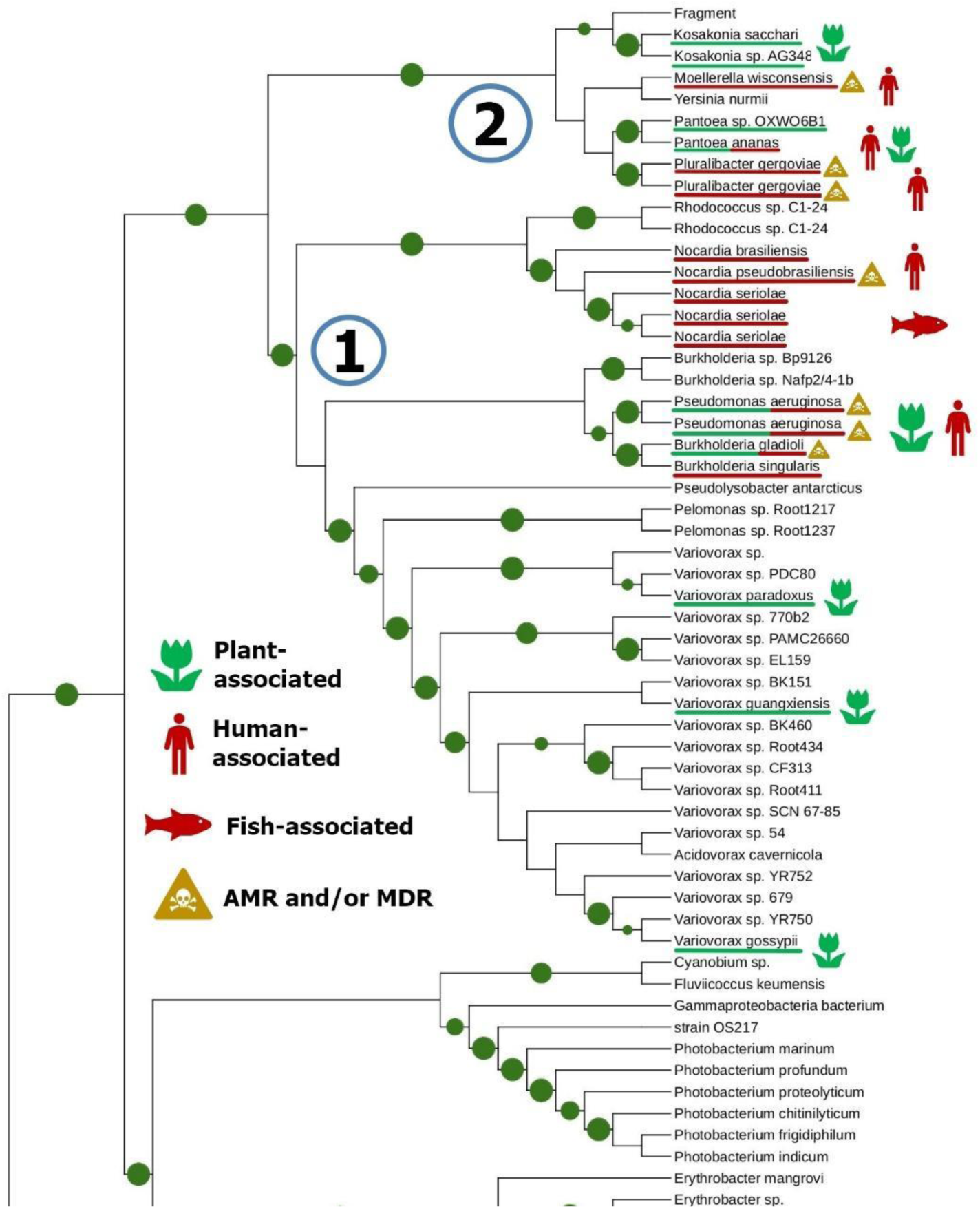
Clusters 1 and 2 of bacterial LOXs reflect two independent series of horizontal gene transfers between plant symbionts, versatile pathogens, and vertebrate/human pathogens. Here, only a fragment of the phylogenetic tree is represented (the full tree is depicted in the **Figure S1**). The circle diameters depict the bootstrap values (display range of 0.5 to 1.0). *Created with iTOL*

In contrast, the LOXs of bacteria associated with marine invertebrates were predominantly separated to the distinct cluster (we call it “Cluster 4”, **Figure 4**, **Figure S2**) and a distinct subcluster within the “Cluster 3” along with plant and human/vertebrate pathogens. Notably, no versatile (cross-kingdom) pathogens were included in the Cluster 3 (**Figure 5**, **Figure S3**), and it is the only cluster where the possible “terrestrial-marine” LOX gene transfer was observed. So, this cluster is significantly dissimilar from the Clusters 1 and 2, although this phylogenetic picture still corresponds to the collective ecological profile described above in terms of low connectivity between “marine-related” terms and “plant-related”/“human-related” term groups.

**Figure 4.**
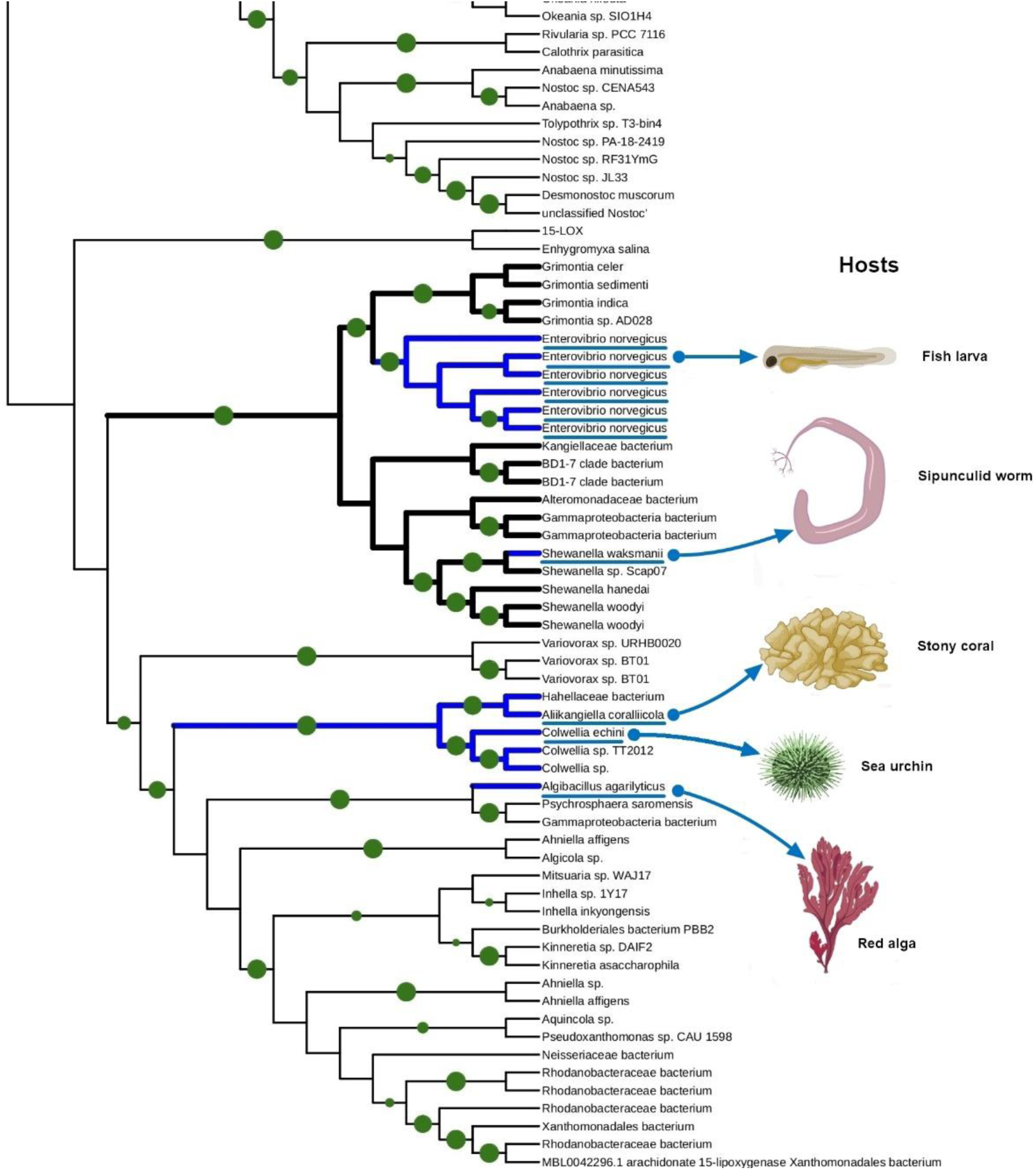
The fragment of phylogenetic tree depicting the “Cluster 4” of bacterial LOXs. This cluster encompasses bacteria isolated from different aquatic organisms: a red alga, a sea urchin, a stony coral, a sipunculid worm, and a fish larva. The circle diameters depict the bootstrap values (display range of 0.7 to 1.0). *Created with iTOL and BioRender.com*

**Figure 5.**
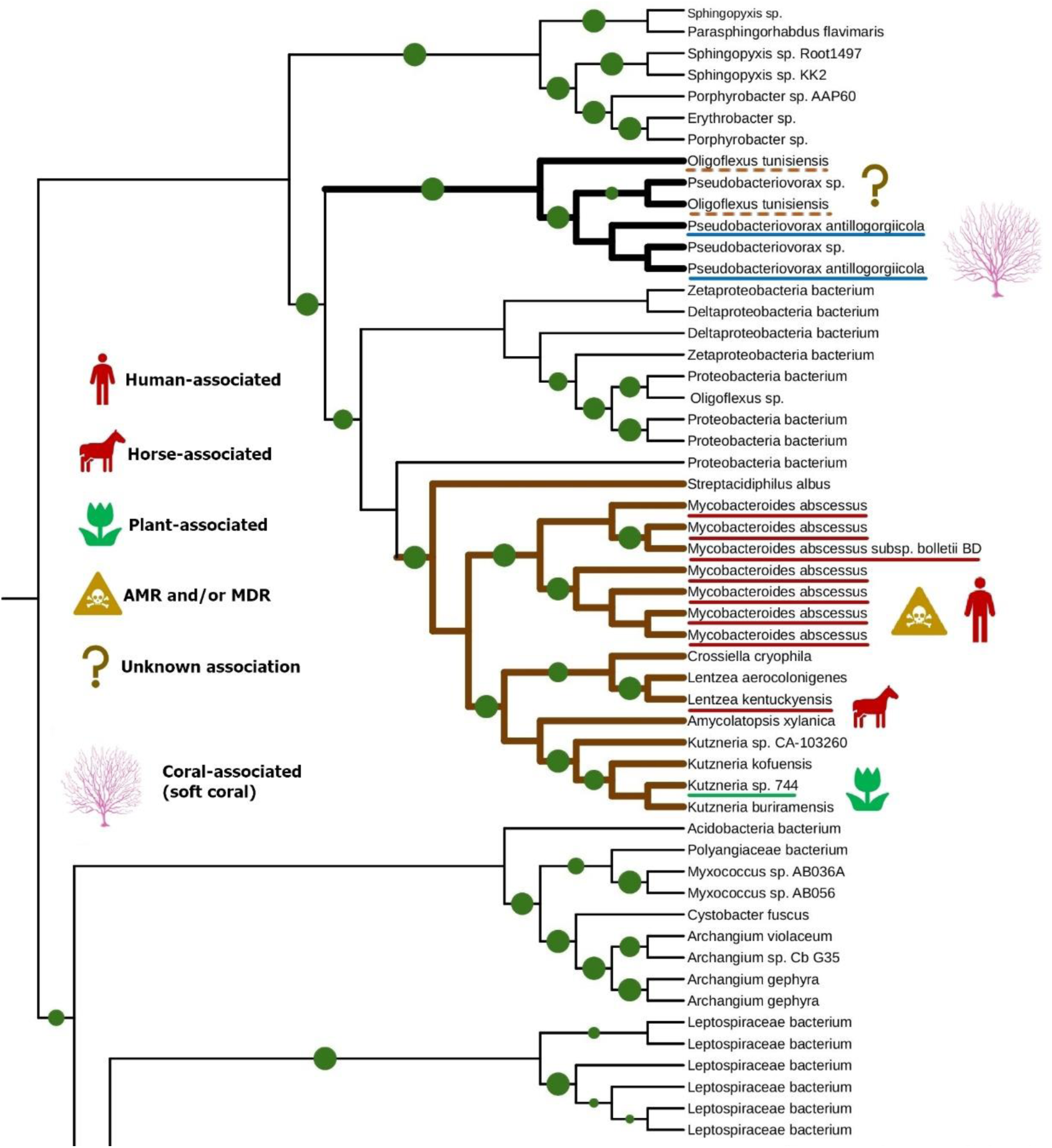
This fragment of a phylogenetic tree depicts the “Cluster 3” of bacteria LOXs — the only cluster, where the transfer between plant-and vertebrate associated bacteria and a coral-associated bacteria was recorded. The circle diameters depict the bootstrap values (display range of 0.7 to 1.0). *Created with iTOL and BioRender.com*

Regarding the phylogenetic relationship of these four clusters, our data are only partially conclusive due to the insufficient number of gap-free sites while analyzing the whole dataset. However, we constructed the phylogenetic network including subsets of Clusters 1, 2, and 3, which remained after applying MaxAlign procedure to make the number of gap-free-sites sufficient. This network also included outgroups, such a cyanobacteria and myxobacteria. To construct this phylogenetic network, we used the same sequence subset as for the phylogenetic tree depicting Cluster 1 and 2. It did not include Cluster 4 due to significant distance with it — it was included in another subset. However, even at the network construction stage, we had indirect evidence that the phylogenetic distance between Clusters 1, 2, and 3 is significantly less than the phylogenetic distance between each of these clusters and Cluster 4.

The resulting network showed that Clusters 1, 2, and 3 were located closely to each other and formed a kind of “supercluster” (**Fig. 6**). Within this study, this was the first evidence that lipoxygenases of Cluster 3 are closely related to lipoxygenases of Clusters 1 and 2.

**Figure 6.**
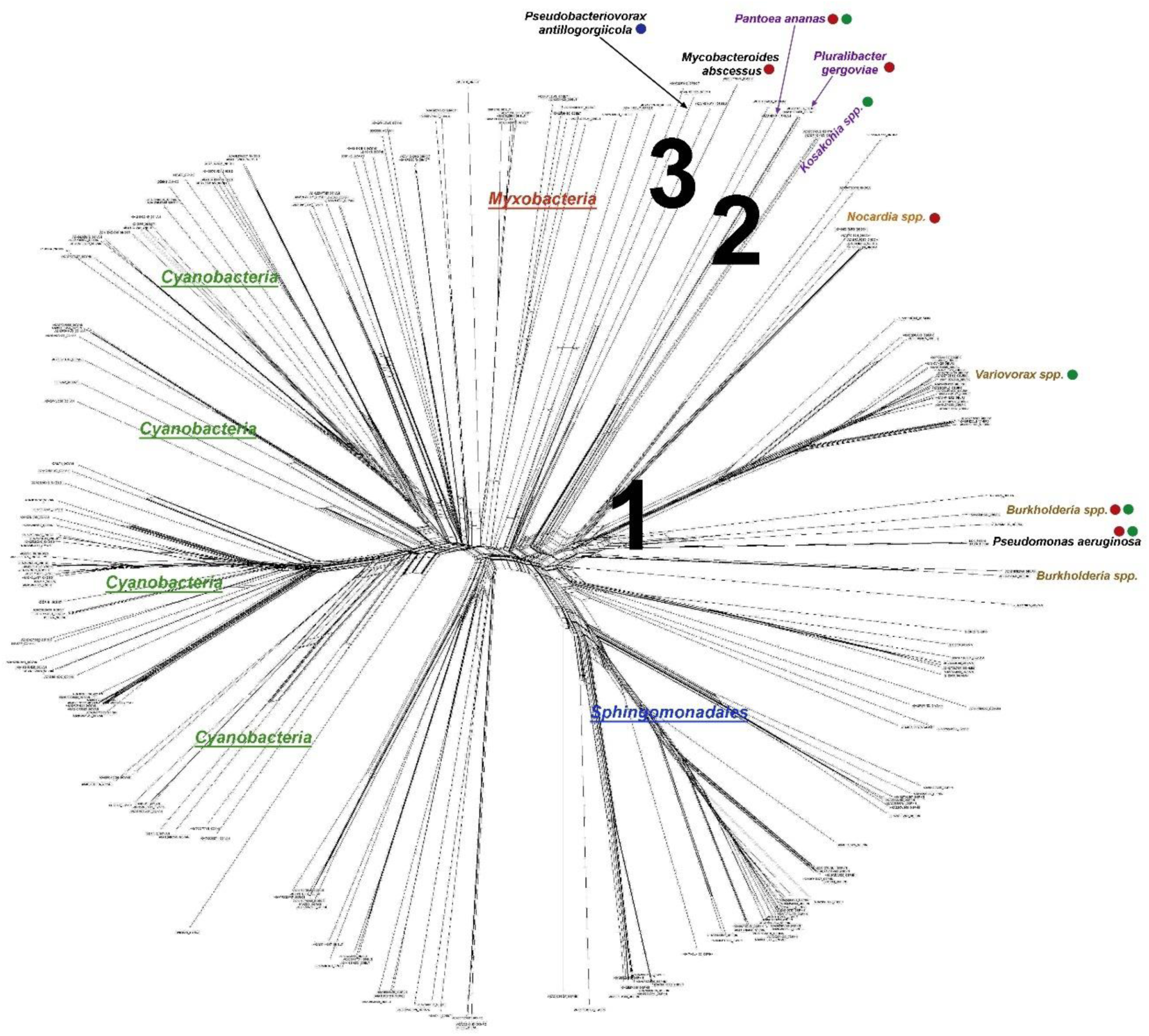
SplitsTree phylogenetic network for lipoxygenases of Clusters 1, 2, and 3 shows their close location. Cyanobacteria, Myxobacteria, and Sphingomonadales were added as large outgroups. The cluster numbering and the color coding of ecological functions correspond to those in those in the Figures 3 and 5. *Created with SplitsTree*.

The “supercluster” (comprising Clusters 1, 2, 3) and Cluster 4 seem to be located distantly, and this result corroborates our earlier results and results of other researchers. In our previous paper, this “supercluster” is also evident in the phylogenetic network of the larger dataset of bacterial and eukaryotic LOXs. However, due to larger dataset and more diverse sequences, MaxAlign procedure excluded more sequences, including all Cluster 2. But Cluster 1 and Cluster 3 are still located closely together in the phylogenetic network in our previous paper (**Figure 7**).

**Figure 7.**
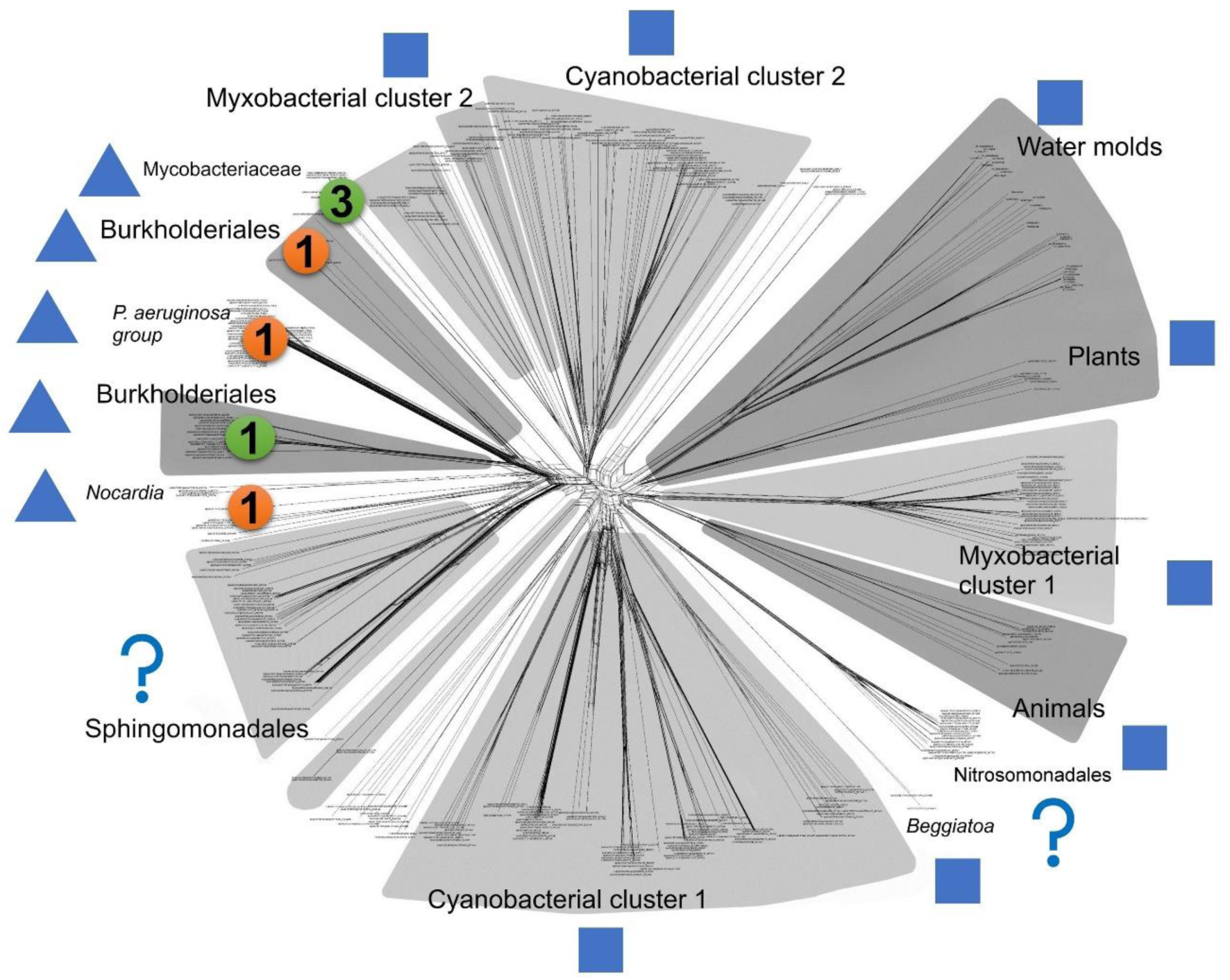
*SplitsTree* phylogenetic network from our previous paper [8] with markers added. Squares depict multicellular taxa, and triangles depict taxa involved in any kind of host-microbe interaction. It is evident that the diagram is divided by these two ecological functions. However, there are taxa for which multicellularity or host-microbe interactions have not been described; they are marked with question mark. All taxa marked with triangle cluster closely together despite they include the representatives of “Cluster 1” (the number “1” in a round) and “Cluster 3” (the number “3” in a round). This relationship is similar to the one presented in the Fig. 6. This suggests that the Clusters 1, 2, and 3 are closely related. *Image credit: Kurakin G. et al. (2020)* [8]*, amended*.

These results corroborate the aforementioned fact that lipoxygenases of “marine-related” bacteria are phylogenetically distant from their counterparts in “plant-related” and “human-related” bacteria, and these later groups, in turn, are phylogenetically close to each other. Taken together, these data indicate that the bacterial LOX phylogeny is associated with host-microbe interactions in several independent clusters. This means that LOXs probably faced host-associated selection pressure. In turn, this means that bacterial LOXs are biochemically involved in host-microbe interaction in different host types: plants, vertebrates, and invertebrates.

### Binding site analysis reveals that (ω-6)S-specificity of bacterial LOXs may contribute to plant-human host jumps

Our analysis also suggests that in the course of evolution, bacterial LOXs might swiftly switch their function from plant-microbe interaction to human-microbe interaction and *vice versa*. Moreover, such switches can ever occur in the same species without significant evolution of the LOX sequence — it is the case of versatile pathogens capable of cross-kingdom host jumps. Conversely, function switches between plant-or human-microbe interaction and interaction with marine hosts are significantly less common. It could be explained by the environmental difference: both plants and humans live in the terrestrial environment and have much more chances to contact each other than to contact any organism from the aquatic environment. However, we decided to check if any biochemical properties of these LOXs determine and explain their ecology.

We used ConSurf and additional statistical analysis to find any differences in key residues determining the substrate insertion direction, stereospecificity, and regiospecificity. These residues had been already characterized in papers [148, 149]. We have not found any significant differences between studied bacterial lipoxygenases in the terms of insertion-determining residues (probably the “tail first” insertion in all lipoxygenases) and Coffa residue (**Ala** in all our LOXs, conservation score = 9, which means that all LOXs in our dataset have S-stereospecificity).

LOX regiospecificity is determined by the bottom triad of the substrate-binding site — 3 amino acid residues forming the binding site bottom. We calculated the total volume of these residues (as the sum of residue volumes) and performed Mann-Whitney test for three groups of LOXs:

1. LOXs of human-and vertebrate-associated bacteria (n=12);
2. LOXs of plant-associated bacteria (n=10);
3. LOXs of marine-associated bacteria (n=6)

(the LOXs of cross-kingdom pathogens were included both in the “plant” and the “human-vertebrate” samples).

There was no statistically significant difference between the “human/vertebrate group” and the “plant group” (U=31). But there were statistically significant differences between each of the above groups and the “marine group” (U=6 and U=2, respectively) even when checking the null hypothesis against p=0.01 (it approximately corresponds to p=0.05 with Bonferroni correction).

Indeed, the LOXs of human-associated bacteria and the LOXs of plant-associated bacteria have on average almost the same total volumes of the binding site bottom triad (mean=475.24×10^-3^ nm^3^, σ=30.45×10^-3^ nm^3^ and mean=497.27×10^-3^ nm^3^, σ=30.82×10^-3^ nm^3^, respectively). The bottom volumes of the LOXs of marine-associated bacteria were much lower (mean=411.87×10^-3^ nm^3^, σ=21.72×10^-3^ nm^3^). This is graphically shown in the box plot (**Figure 8**). The data for all bacteria included in this statistical analysis are provided in **Table S4**.

**Figure 8.**
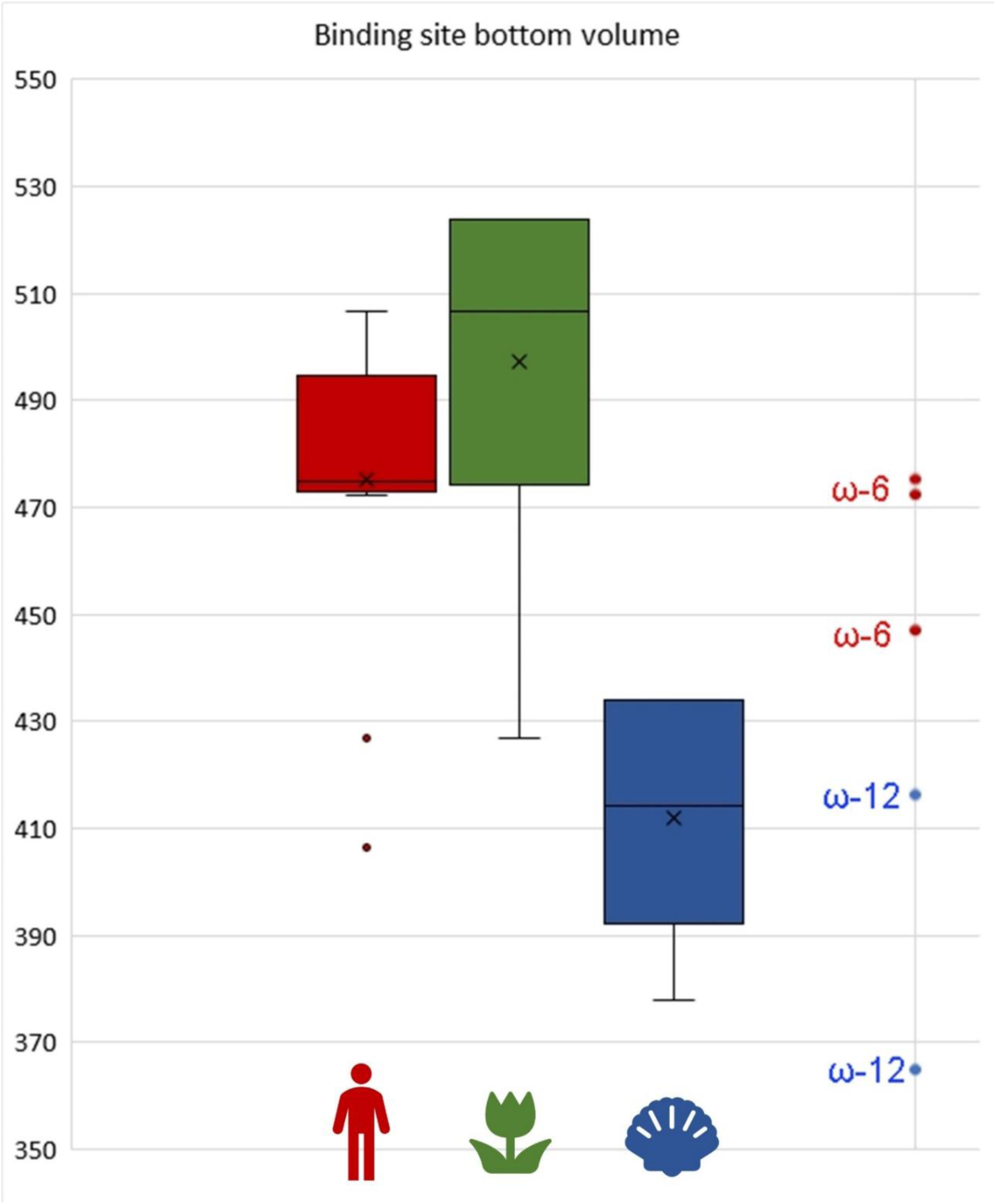
The box plot of total LOX ligand binding site bottom volumes for “human/vertebrate-associated” (**red**), “plant-associated” (**green**), and “marine-associated” (**blue**) groups. All volumes are given in 10^-3^ nm^3^ according to Perkins (1986) [146]. LOXs of “human/vertebrate-associated” and “plant-associated” groups have total bottom volumes at almost the same level; in contrast, LOXs of the “marine-associated” group have significantly lower bottom volumes (p<0.05 with Bonferroni correction). The bar at the right shows total binding site bottom volumes of experimentally characterized LOXs [149, 151–156] along with their activities. “Human-related” and “plant-related” groups correspond (ω-6)-LOX activity while “marine-related” group is shifted towards (ω-12)-LOX activity.

The box plot (**Figure 8**) also graphically shows the presence of two outliers in the “human-vertebrate” group with extremely low total bottom volume. But they fully correspond to the average values for the “marine” group and lie within 1σ from its mean. These LOXs phylogenetically belong to the Cluster 3 — the only cluster where the ancient terrestrial-marine LOX gene transfer was recorded.

Compared with the binding site bottom volumes of experimentally characterized LOXs [148, 150–155] (calculated by the same method), the “human-related” and “plant-related” LOX groups appear to correspond to (ω-6)-LOXs. The *Pseudomonas aeruginosa* and *Burkholderia thailandensis* LOXs (both belonging to the “human/vertebrate” group in our statistics) have total binding site bottom volumes (472.2×10^-3^ nm^3^ and 475×10^-3^ nm^3^, respectively) near to the mean of the “human-related” group. The mean of the “plant-related” group is even more. It must also correspond to the (ω-6)S-LOX activity because it requires the minimally profound substrate penetration into the active site (and, thus, the maximal volume of the site bottom). So, we may conclude that (ω-6)S-LOX activity is required for plant-vertebrate host jumps. Conversely, the “marine-related” group is significantly shifted towards (ω-12)-LOXs. It means that different ecological groups of bacterial LOXs statistically correspond to different biochemical activities.

## Discussion

### Possible mechanism of the LOX-mediated plant-human host jumps

We have for the first time confirmed that pathogen and symbiont LOXs are associated with a broad host range and cross-kingdom host jump ability. The correspondence between the LOX phylogeny and the host-microbe interactions also confirms that these enzymes are involved in pathogenesis and symbiosis and probably interact with a host.

One of the pathogen LOXs in our dataset – *Pseudomonas aeruginosa* LoxA — was earlier experimentally characterized providing the pathophysiological insights into this interaction [150, 156, 157]. This LOX is represented in our dataset, our statistical analysis sample and in the Cluster 1 of the phylogenetic tree and attributed to be a LOX of a cross-kingdom pathogen because of a broad host range of *Pseudomonas aeruginosa*. However, the available experimental data regard only the role of *Pseudomonas aeruginosa* LOX in interactions with a human organism.

In addition to its direct destructive action on invaded tissues [156, 157], this LOX is involved in the complicated signalling crosstalk with human leukocytes. When invading human tissues, this pathogen secretes a 15S-LOX which converts the host’s arachidonic acid to 15S-ΗΕΤΕ which is further metabolized by the human leukocytes to anti-inflammatory mediators such as lipoxin A_4_ (**Figure 9**). Their action, in turn, leads to decrease of immune cell recruiting and immune response itself, and, finally, to the invasion facilitation [150].

**Figure 9.**
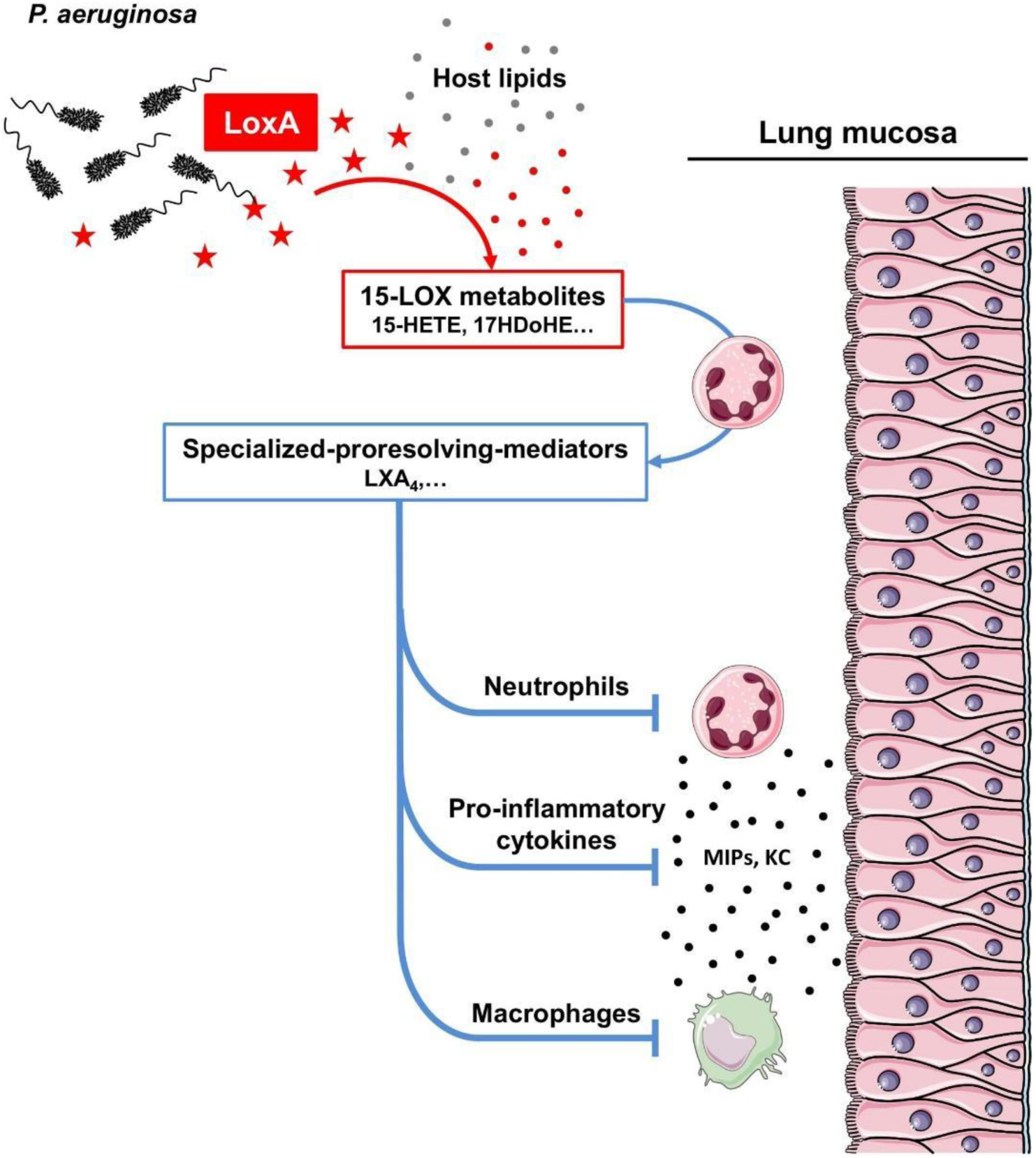
The graphical outline of *Pseudomonas aeruginosa* LOX action to facilitate a host’s tissue invasion. ***Image credit:*** Morello, E. et al. (2019) *Frontiers in Microbiology* [150], CC BY 4.0

Host-pathogen oxylipin crosstalk is not restricted to humans and is widely exploited by bacterial and fungal plant pathogens, but the exact biochemical mechanisms differ case by case [158]. Our results suggest that some bacteria share similar LOXs and biochemical pathways for interacting with different hosts — from plants to corals.

Moreover, we could now assume that both plant pathogenesis and plant symbiosis may be driven by similar bacterial LOXs. This is the finding of particular interest because plant-symbiont oxylipin signalling remains poorly elucidated: one of the most comprehensive reviews on this topic by Beccacioli et al. [158] cites only one example of such interaction — the paper on beneficial interaction between the fungus *Trichoderma virens* and maize (*Zea mays*) [159]. No instances of oxylipin-mediated crosstalk between plants and their bacterial symbionts were cited. Our results highlight the possible way of such oxylipin signalling.

Our data also show that LOX genes can make horizontal transfers thus finding new ecological niche for themselves (like Richard Dawkins’s “selfish genes” [160]) and enable their carriers to make cross-kingdom host jumps. Such species are discussed above as versatile pathogens.

According to our data, versatile LOX-carrying bacteria affect plants, vertebrates, sometimes insects, but never affect marine invertebrates. Furthermore, LOX transfers between plant-associated and vertebrate-associated pathogens are the most frequent events according to our phylogeny (at least 3 independent series of plant-human transfers), while LOX transfers between plant-or human-associated and marine-associated pathogens are rarer (only one even according to our phylogeny). In our ecological networks, a small “marine-related” cluster is located apart from the “plant-related” and “human-related” clusters. It leads to an assumption that bacterial lipoxygenases can easily switch from a plant host to a vertebrate host (and *vice versa*), but a switch to a marine host is very uncommon. The simplest explanation could be the ecological isolation: both plants and humans live in the terrestrial environment and have much more tight contact between each other than with any aquatic organism. According to this hypothesis, if humans were aquatic mammals (like dolphins) and lived in a coral reef ecosystem, we could observe multiple “coral-human” host jumps. This hypothesis assumes full “compatibility” of pathogen/symbiont LOXs with any organism: vertebrate, insect, plant, or marine invertebrate.

However, we have found that LOXs themselves have structural differences in the ligand-binding site which are linked to the host type. The “plant-compatible” and the “vertebrate-compatible” LOXs appeared to have no statistically significant differences in the binding site structure in contrast to the “marine-compatible” LOXs. It leads to an assumption that bacterial LOXs face some biochemical “compatibility requirements” for successful host-microbe interactions. According to this hypothesis, bacterial LOXs are capable of cross-kingdom switches because plant and animal hosts require the same regio-and stereospecificity.

Matching statistical data on LOX binding site bottom volumes to experimental data for biochemically characterized bacterial LOXs enabled us to infer the LOX specificity needed both for colonizing plants and humans. It must correspond to the (ω-6)S-LOX activity.

In the case of human (and other vertebrates), any (ω-6)S-LOX of a pathogen is expected to behave like *Pseudomonas aeruginosa* LOX: for arachidonic acid prevalent in the vertebrate pool of polyunsaturated fatty acids (PUFAs), (ω-6)S-LOX activity means 15S-LOX activity, which contributes to the lipoxin biosynthetic pathway and suppresses the host’s immune response (which is fulfilled in the case of *Pseudomonas aeruginosa*).

But what about plants? In contrast to humans, we have no direct experimental data on the interaction of any bacterial LOX with them. However, we could biochemically infer this interaction. In the case of linolenic acid (abundant in plant cells), any (ω-6)S-LOX could be predicted to produce 13S-HpOT, which is a normal precursor of jasmonates. So, if (ω-6)S-LOX enhances lipoxin production in the human tissues, they should enhance jasmonate production in the plant body.

Here, the “explanative gap” is fully closed because jasmonate signalling hijacking is a well-known trick of some plant pathogens. They increase a plant’s susceptibility by inappropriate activation of jasmonate signalling pathway. However, almost all instances of such pathogenesis known up do date involve the use of toxin mimicking the natural jasmonate or acting as prohormone. The best-known example is *Pseudomonas syringae* that uses coronatine activating the jasmonate receptor JAZ-COI1 [161] to facilitate the invasion. A grapevine pathogen *Lasiodiplodia mediterranea* facilitates its invasion by the prohormone toxin lasiojasmonate A [162]. The current study provides computational evidence that bacteria might use lipoxygenase to “spoof” the plant immunity with natural jasmonates rather than jasmonate-mimicking toxin. By accident or by the convergent evolution, the same LOX activity appeared to be “compatible” with a human lipoxin pathway which enabled LOX-carrying bacteria to be versatile pathogens (**Figure 10**).

**Figure 10.**
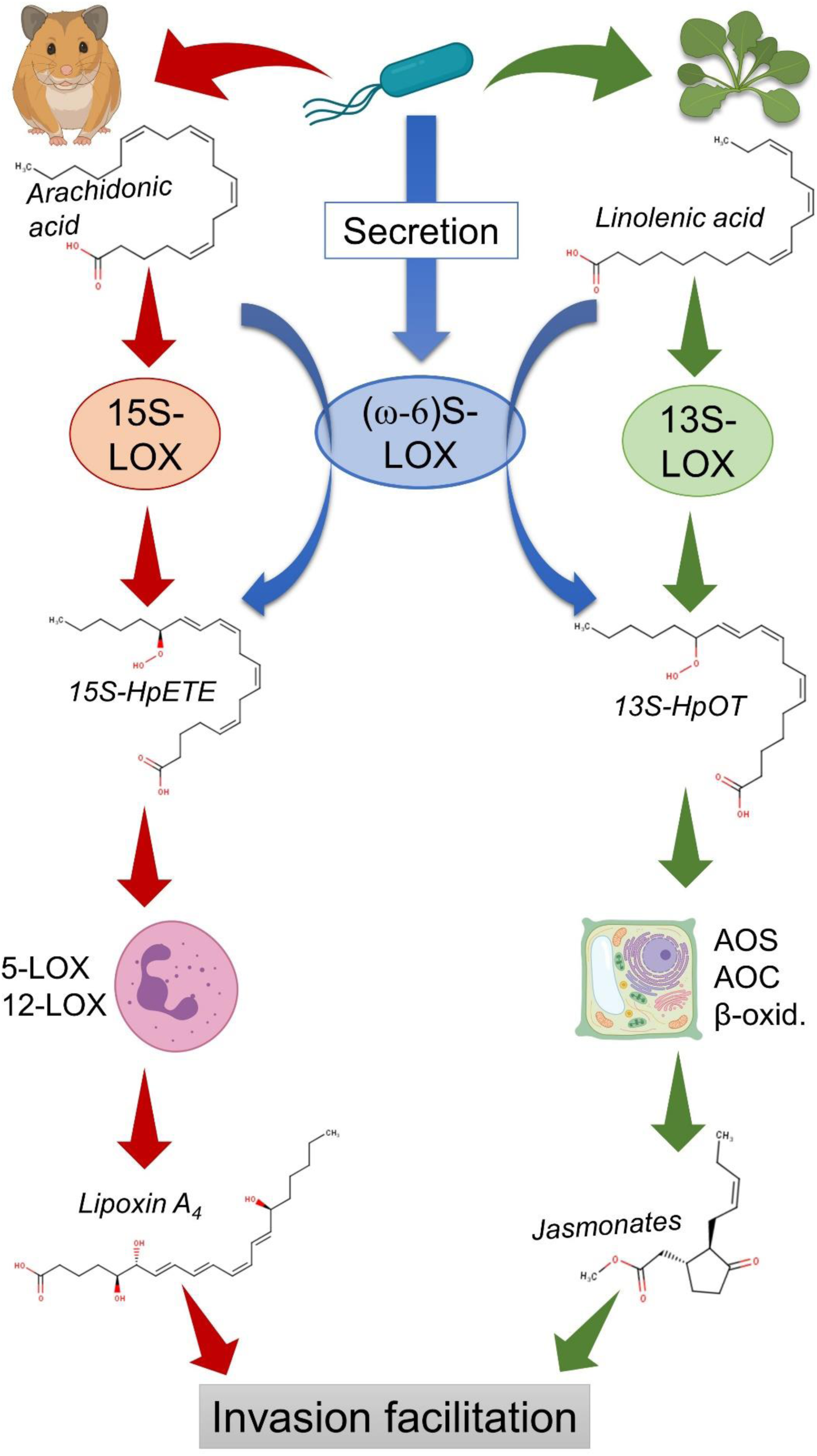
Hypothetical mechanism of plant-animal host jumps provided by bacteria LOXs. In humans and vertebrates, a secreted bacterial (ω-6)S-LOX oxidizes arachidonic acid to form 15S-HpETE, which is further converted into lipoxins in leukocytes by other LOX isozymes. Lipoxins suppress inflammation and immune response thus facilitating the invasion. *Created with BioRender.com*

The immune response suppression hypothesis partially explains why this feature is shared by plant pathogens and plant symbionts (as well as insect pathogens and symbionts): the both groups need to evade the host’s immune response to survive in its body. However, this raises an additional question: why do human symbionts not use this strategy?

It is not the only specific thing of interaction between LOX-carrying bacteria and a human (or a vertebrate) organism. As discussed above, the ability of such pathogens to affect humans is extremely strongly associated with nosocomial or opportunistic traits. It means that these bacteria are prone to affect immunocompromised people (the terms “compromised” and “immunocompromised” are also present in the “human-related” cluster, but have relatively low connectivity compared to hubs described above). Conversely, in the case of plants and insects, we did not observe such associations: LOX carriers are capable of full-fledged pathogenesis and symbiosis.

These differences are tightly connected: if a pathogen oxylipin “spoofing” is insufficient to suppress the healthy human’s immune response, there is no reason to expect that it will be sufficient for human symbionts to evade the immune response and survive. Evidently, there is something in the human (or, more broadly, vertebrate) immune system that makes it more resilient to oxylipin signalling hijacks. Here, we come on shaky grounds: we have insufficient data to explain what it could be. But it is reasonable to suggest that it is the presence of adaptive immunity. Indeed, one of the mechanisms used by human microbiota to evade immune response is immune tolerance — the acquired silencing of a host’s adaptive immunity to the bacterium’s antigens [163].

### Public health risks and molecular epidemiology surveillance

First of all, we should downgrade the alert regarding possible “superspreading” of lipoxygenases between nosocomial bacteria which we postulated in our previous work [8]. The “*Pseudomonas aeruginosa* group” which comprised *Klebsiella pneumoniae*, *Enterobacter cloacae*, *Acinetobacter baumannii,* and other bacteria putatively linked by fast horizontal transfers of LOX genes proved to be an artifact. All LOX sequences of this group excepting *Pseudomonas aeruginosa* sequences were subsequently removed from the databases due to probable contamination. Here, we have found new evidence for the role of horizontal LOX gene transfers in pathogenicity evolution, but we suggest that these processes occur on longer timescales.

The fact that the terms “emerging”, “AMR”, and “MDR” are prevalent in the collective ecological profile of LOX carriers and serve as hubs in the term network representing this profile should raise concerns. Moreover, antimicrobial-resistant and multi-drug resistant bacteria are represented in each of 3 phylogenetic clusters of the pathogen and symbiont LOXs. It means that bacterial LOXs are associated with emerging status, antimicrobial resistance and some other public health threats. Furthermore, multiple connections between these traits and plant or human pathogenicity entitle us to say that emerging pathogens use oxylipin signalling for plant-human (or human-plant) host jumps.

But this conclusion should be considered carefully. Firstly, we have mentioned above that the traits “AMR” and “MDR” could have overestimated weights due to normalization bias. We are sure that this possible bias has limited effect because the used term “AMR” corresponds to the real trait of studied bacteria. Moreover, the term “emerging” has also a very big weight, but cannot be subjected to normalization bias. But on the visual level, it could produce a formidable picture and lead to the emergence of news about “new deadly pathogens” or a “new global epidemic”. We caution all science journalists reading this article against doomsaying.

Secondly, our data provide strong evidence that LOX-carrying pathogens have low epidemic and pandemic potential. As we discussed above, they are prone to affect immunocompromised people or people with specific diseases (e.g., cystic fibrosis), and cannot fully overcome the immune barrier of a healthy human despite multiple recorded host jump events. So, we find no reason to expect a full-fledged epidemic or pandemic caused by LOX-carrying pathogens.

However, all statistical signs of “public health threat” are connected with the extreme danger they pose to specific populations — immunocompromised people or people with comorbidities, especially with cystic fibrosis. Regarding these populations, we cannot rule out the possibility of limited outbreaks. Sporadic cases are also dangerous for these people and could lead to higher mortality and decrease in the quality of life. Thus, bacterial lipoxygenases could be a useful molecular epidemiology marker for oncology, hematology, transplantology, and cystic fibrosis medicine.

We continue updating our “pathogen blacklist” — the list of LOX-carriyng pathogens that can pose danger to humans — on the grounds of publicly available data. The last update before this article was published in English in the *Nature Portfolio Microbiology Community* blog [164] and in Russian in *Priroda* [9]. In this work, this list is incorporated into **Table 1**. But the full-fledged molecular epidemiology surveillance requires much more strain-dependent data. We are open for further collaboration for this purpose.

### Possible implications for marine biology

In contrast to plants and humans, our knowledge about LOX-carrying bacteria in marine organisms is limited. In most cases, we suggested the association with marine organisms only on the basis of isolation of respective bacteria from a particular organism. So, the ecological functions of LOX-carrying bacteria (pathogen/symbiont) are unknown for all “marine-associated” bacteria in our dataset. But we can suggest that in the underwater world, more intriguing host jumps of LOX-carrying bacteria occur. They involve corals, sea urchins, algae, sipunculids, and fishes. We have found that these host jumps require other LOX regiospecificity, than for plants and humans. We cannot infer any mechanism of their actions because of scarce data on oxylipin signalling in marine invertebrates, but can suggest that bacterial LOXs could be a useful tool to study it.

## Conclusions

In this article, we have proposed a mechanism of cross-kingdom host jumps by interacting with evolutionary distant immune systems. The analysis of available data shows that some bacteria may exploit the hosts’ oxylipin signalling systems which are widespread across different kingdoms. The most prevalent are plant-human host jumps because of a biochemical “intercompatibility” of plant and human LOX activities needed for immune response suppression. This finding could be interesting for basic scientists in a wide range of areas — from immunology to marine biology. But it could also be useful for medical microbiology and public health. Bacterial LOXs in a nosocomial environment could indicate an emerging pathogen which could be dangerous for some vulnerable groups of patients. Bacterial LOX carriers isolated from patients may require further consideration in terms of antimicrobial therapy and the particular role of this bacteria in the patient’s condition pathogenesis (as is the case of cystic fibrosis). It is evident that pathogen and symbiont LOXs require further investigation in both basic and applied life sciences.

## Acknowledgements

The author acknowledges:

• Anastasiya Kuznetsova (an independent data visualization expert, https://nastengraph.medium.com/) for her valuable advice in the course of this research;

• Alina Lebedeva (Tver State University, Teaching Assistant at the Department of Language Theory, Translation and French Philology) and Tatyana Shcherbakova (Translator-coordinator, Orient Translating LLC, Saint Petersburg) for literary editing this text;

• Livia Leoni (Roma Tre University, Associate Professor at the Department of Science) for her valuable comments on the correction of this paper;

• Artem Tishkov (I.P. Pavlov Saint Petersburg State Medical University, Head of Physics, Mathematics and Informatics Department) for his valuable advices regarding statistical methods;

• Mikhail Gelfand and Yulia Sarana (Skolkovo Institute of Science and Technology, Center of Molecular and Cellular Biology) for their valuable criticism regarding vibrios.

## Ethics statement

This research did not involve any experimental animals or humans as a research object.

## Data availability

The unedited versions of phylogenetic trees and the full versions of tables are included in this paper as **Supplementary Figures** and **Supplementary Tables**, respectively. Datasets, spreadsheets, interactive data and any other data generated for this research are available by request to the author’s email.

## References

1. Pulliam, J. R. (2008) Viral host jumps: moving toward a predictive framework, EcoHealth, 5, 80–91, doi: 10.1007/s10393-007-0149-6.

2. Letko, M., Seifert, S. N., Olival, K. J., Plowright, R. K., and Munster, V. J. (2020) Bat-borne virus diversity, spillover and emergence, Nat. Rev. Microbiol., 18, 461–471, doi: 10.1038/s41579-020-0394-z.

3. Sturm-Ramirez, K. M., Hulse-Post, D. J., Govorkova, E. A., Humberd, J., Seiler, P., Puthavathana, P., Buranathai, C., Nguyen T. D., Chaisingh, A., Long, H. T., Naipospos, T. S. P., Chen, H., Ellis, T. M., Guan, Y., Peiris J. S. M., and Webster, R. G. (2005) Are ducks contributing to the endemicity of highly pathogenic H5N1 influenza virus in Asia?, J. Virol., 79, 11269–11279, doi: 10.1128/JVI.79.17.11269-11279.2005.

4. Taylor, J., and Pelchat, M. (2010) Origin of hepatitis δ virus, Future Microbiol., 5, 393–402, doi: 10.2217/fmb.10.15.

5. Kirzinger, M. W., Nadarasah, G., and Stavrinides, J. (2011) Insights into cross-kingdom plant pathogenic bacteria, Genes, 2, 980–997, doi: 10.3390/genes2040980.

6. Van Baarlen, P., Van Belkum, A., Summerbell, R. C., Crous, P. W., and Thomma, B. P. (2007) Molecular mechanisms of pathogenicity: how do pathogenic microorganisms develop cross-kingdom host jumps?, FEMS Microbiol. Rev., 31, 239–277, doi: 10.1111/j.1574-6976.2007.00065.x.

7. Rahme, L. G., Ausubel, F. M., Cao, H. et al. (2000) Plants and animals share functionally common bacterial virulence factors, Proc. Natl. Acad. Sci. USA, 97(16), 8815–8821, doi: 10.1073/pnas.97.16.8815

8. Kurakin, G. F., Samoukina, A. M., and Potapova, N. A. (2020) Bacterial and Protozoan Lipoxygenases Could be Involved in Cell-to-Cell Signaling and Immune Response Suppression, Biochemistry (Moscow), 85, 1048–1063, doi: 10.1134/S0006297920090059

9. Kurakin., G. F. (2022) Oksilipiny bakterij: kljuch k mnogokletochnosti i bor’be s ustojchivost’ju k antibiotikam? [Bacterial oxylipins: a key to multicellularity and to combating antimicrobial resistance?], Priroda, 2, 26–32, doi: 10.7868/S0032874X2202003X

10. Toukabri, W., Ferchichi, N., Hlel, D., Jadlaoui, M., Kheriji, O., Mhamdi, R., and Trabelsi, D. (2021) Response of intercropped barley and fenugreek to mono-and co-inoculation with *Sinorhizobium meliloti* F42 and *Variovorax paradoxus* F310 under contrasting agroclimatic regions, Arch. Microbiol., 203, 1657–1670, doi: 10.1007/s00203-020-02180-8.

11. Garcia Teijeiro, R., Belimov, A. A., and Dodd I.C. (2020) Microbial inoculum development for ameliorating crop drought stress: A case study of *Variovorax paradoxus* 5C-2, N. Biotechnol., 56, 103–113, doi: 10.1016/j.nbt.2019.12.006.

12. Gao, J. L., Yuan, M., Wang, X. M., Qiu, T. L., Li, J. W., Liu, H. C., Li, X. A., Chen, J., and Sun, J. G. (2015) Variovorax guangxiensis sp. nov., an aerobic, 1-aminocyclopropane-1-carboxylate deaminase producing bacterium isolated from banana rhizosphere, Antonie van Leeuwenhoek, 107, 65–72, doi: 10.1007/s10482-014-0304-3.

13. Kämpfer, P., Busse, H. J., McInroy, J. A., and Glaeser, S. P. (2015) *Variovorax gossypii* sp. nov., isolated from *Gossypium hirsutum*, Int. J. Syst. Evol. Microbiol., 65, 4335–4340, doi: 10.1099/ijsem.0.000581.

14. Belli, G., Giovannini, M., Dolce, D., Terlizzi, V., Orioli, T., and Taccetti, G. (2021) *Burkholderia gladioli* infection in a pediatric patient with cystic fibrosis: the clinical challenges of an emergent pathogen, Minerva Pediatr. (Torino), 73, 468–470, doi: 10.23736/s2724-5276.20.05836-3.

15. Cui, G., Yin, K., Lin, N., Liang, M., Huang, C., Chang, C., Xi, P., and Deng, Y. Z. (2020) *Burkholderia gladioli* CGB10: A Novel Strain Biocontrolling the Sugarcane Smut Disease, Microorganisms, 8, 1943, doi: 10.3390/microorganisms8121943.

16. Bedir Demirdag, T., Ozkaya Parlakay, A., Aygar, I. S., Gulhan B., and Kanik Yuksek, S. (2020) Major Aspects of *Burkholderia gladioli* and *Burkholderia cepacia* Infections in Children, Pediatr. Infecti. Dis. J., 39, 374–378, doi: 10.1097/INF.0000000000002587.

17. Zanotti, C., Munari, S., Brescia, G., and Barion, U. (2019) *Burkholderia gladioli* sinonasal infection, Eur. Ann. Otorhinol. Head Neck Dis., 136, 55–56, doi: 10.1016/j.anorl.2018.01.011

18. Dursun, A., Zenciroglu, A., Karagol, B. S., Hakan, N., Okumus, N., Gol, N., and Tanir, G. (2012) *Burkholderia gladioli* sepsis in newborns, Eur. J. Pediatr., 171, 1503–1509, doi: 10.1007/s00431-012-1756-y.

19. Jones, C., Webster, G., Mullins, A. J., Jenner, M., Bull, M. J., Dashti, Y., Spilker, T., Parkhill, J., Connor, T. R., LiPuma, J. J., Challis, G. L., and Mahenthiralingam, E. (2021) Kill and cure: genomic phylogeny and bioactivity of *Burkholderia gladioli* bacteria capable of pathogenic and beneficial lifestyles, Microb. Genom., 7, mgen000515, doi: 10.1099/mgen.0.000515.

20. Marom, A., Miron, D., Wolach, B., Gavrieli, R., and Rottem, M. (2018) *Burkholderia gladioli*-associated facial pustulosis as a first sign of chronic granulomatous disease in a child-Case report and review, Pediatr. Allergy Immunol., 29, 451–453, doi: 10.1111/pai.12884.

21. Ritterband, D., Shah, M., Cohen, K., Lawrence, J., and Seedor, J. (2002) *Burkholderia gladioli* keratitis associated with consecutive recurrent endophthalmitis, Cornea, 21, 602– 603, doi: 10.1097/00003226-200208000-00014.

22. Boyanton, B. L., Jr, Noroski, L. M., Reddy, H., Dishop, M. K., Hicks, M. J., Versalovic, J., and Moylett, E. H. (2005) *Burkholderia gladioli* osteomyelitis in association with chronic granulomatous disease: case report and review, Pediatr. Infect. Dis. J., 24, 837– 839, doi: 10.1097/01.inf.0000177285.44374.dc.

23. Zhou, F., Ning, H., Chen, F., Wu, W., Chen, A., and Zhang, J. (2015) *Burkholderia gladioli* infection isolated from the blood cultures of newborns in the neonatal intensive care unit, Eur. J. Clin. Microbiol. Infect. Dis., 34, 1533–1537, doi: 10.1007/s10096-015-2382-1.

24. Vandamme, P., Peeters, C., De Smet, B., Price, E. P., Sarovich, D. S., Henry, D. A., Hird, T. J., Zlosnik, J., Mayo, M., Warner, J., Baker, A., Currie, B. J., and Carlier, A. (2017) Comparative Genomics of *Burkholderia singularis* sp. nov., a Low G+C Content, Free-Living Bacterium That Defies Taxonomic Dissection of the Genus *Burkholderia*, Front. Microbiol., 8, 1679, doi: 10.3389/fmicb.2017.01679.

25. Wang, Y., Hoffmann, J. P., Chou, C. W., Höner Zu Bentrup, K., Fuselier, J. A., Bitoun, J. P., Wimley W. C., and Morici, L.A. (2020) *Burkholderia thailandensis* outer membrane vesicles exert antimicrobial activity against drug-resistant and competitor microbial species, J. Microbiol., 58, 550–562, doi: 10.1007/s12275-020-0028-1.

26. Klaus, J. R., Majerczyk, C., Moon, S., Eppler, N. A., Smith, S., Tuma, E., Groleau, M. C., Asfahl, K. L., Smalley, N. E., Hayden, H. S., Piochon, M., Ball, P., Dandekar, A. A., Gauthier, C., Déziel, E., and Chandler, J. R. (2020) *Burkholderia thailandensis* Methylated Hydroxyalkylquinolines: Biosynthesis and Antimicrobial Activity in Cocultures, Appl. Environ. Microbiol., 86, e01452–20, doi: 10.1128/AEM.01452-20

27. Garcia E. C. (2017) *Burkholderia thailandensis*: Genetic Manipulation, Curr. Protoc. Microbiol., 45, 4C.2.1–4C.2.15, doi: 10.1002/cpmc.27

28. Vitale, A., Paszti, S., Takahashi, K., Toyofuku, M., Pessi, G., and Eberl, L. (2020) Mapping of the Denitrification Pathway in *Burkholderia thailandensis* by Genome-Wide Mutant Profiling, J. Bacteriol., 202, e00304–20, doi: 10.1128/JB.00304-20

29. Place, D. E., Briard, B., Samir, P., Karki, R., Bhattacharya, A., Guy, C. S., Peters, J. L., Frase, S., Vogel, P., Neale, G., Yamamoto, M., and Kanneganti, T. D. (2020) Interferon inducible GBPs restrict *Burkholderia thailandensis* motility induced cell-cell fusion, PLoS Pathog., 16, e1008364, doi: 10.1371/journal.ppat.1008364

30. Pilátová, M. and Dionne, M. S. (2012) *Burkholderia thailandensis* is virulent in *Drosophila melanogaster*, PloS One, 7, e49745, doi: 10.1371/journal.pone.0049745

31. Garcia, E. C. and Cotter, P. A. (2016) *Burkholderia thailandensis*: Growth and Laboratory Maintenance, Curr. Protoc. Microbiol., 42, 4C.1.1–4C.1.7, doi: 10.1002/cpmc.15

32. Lei, X., Zhao, R., Geng, Y., Wang, K., Yang, P. O., Chen, D., Huang, X., Zuo, Z., He, C., Chen, Z., Huang, C., Guo, H., and Lai, W. (2020) *Nocardia seriolae*: a serious threat to the largemouth bass *Micropterus salmoides* industry in Southwest China, Dis. Aquat. Organ., 142, 13–21, doi: 10.3354/dao03517

33. Han, H. J., Kwak, M. J., Ha, S. M., Yang, S. J., Kim, J. D., Cho, K. H., Kim, T. W., Cho, M. Y., Kim, B. Y., Jung, S. H., and Chun, J. (2019) Genomic characterization of *Nocardia seriolae* strains isolated from diseased fish, MicrobiologyOpen, 8, e00656, doi: 10.1002/mbo3.656

34. Kim, J. D., Lee, N. S., Do, J. W., Kim, M. S., Seo, H. G., Cho, M., Jung, S. H., and Han, H. J. (2018) *Nocardia seriolae* infection in the cultured eel *Anguilla japonica* in Korea, J. Fish Dis., 41, 1745–1750, doi: 10.1111/jfd.12881

35. Wang, P. C., Chen, S. D., Tsai, M. A., Weng, Y. J., Chu, S. Y., Chern, R. S., and Chen, S. C. (2009) *Nocardia seriolae* infection in the three striped tigerfish, *Terapon jarbua* (Forsskål), J. Fish Dis., 32, 301–310, doi: 10.1111/j.1365-2761.2008.00991.x

36. Hou, S., Wang, W., Chen, G., Xia, L., Wang, Z., and Lu, Y. (2021) Identification of a secreted superoxide dismutase (SOD) from *Nocardia seriolae* which induces apoptosis in fathead minnow (FHM) cells, J. Fish Dis., 44, 63–72, doi: 10.1111/jfd.13268

37. Hou, S., Chen, G., Wang, W., Xia, L., Wang, Z., and Lu, Y. (2020) Identification of a cell-wall peptidase (NlpC/P60) from *Nocardia seriolae* which induces apoptosis in fathead minnow cells, J. Fish Dis., 43, 571–581, doi: 10.1111/jfd.13154

38. Yasuike, M., Nishiki, I., Iwasaki, Y., Nakamura, Y., Fujiwara, A., Shimahara, Y., Kamaishi, T., Yoshida, T., Nagai, S., Kobayashi, T., and Katoh, M. (2017) Analysis of the complete genome sequence of *Nocardia seriolae* UTF1, the causative agent of fish nocardiosis: The first reference genome sequence of the fish pathogenic Nocardia species, PloS One, 12, e0173198, doi: 10.1371/journal.pone.0173198

39. Makadia, S., Patel, I., Soosaipillai, I., and Tarasiuk-Rusek, A. (2020) First Case of *Nocardia pseudobrasiliensis* Causing Primary Cutaneous Nocardiosis in an Immunocompetent Patient, J. Investig. Med. High Impact Case Rep., 8, 2324709620938228, doi: 10.1177/2324709620938228

40. Ruimy, R., Riegel, P., Carlotti, A., Boiron, P., Bernardin, G., Monteil, H., Wallace, R. J., Jr, and Christen, R. (1996) *Nocardia pseudobrasiliensis* sp. nov., a new species of Nocardia which groups bacterial strains previously identified as *Nocardia brasiliensis* and associated with invasive diseases, Int. J. Syst. Bacteriol., 46, 259–264, doi: 10.1099/00207713-46-1-259

41. Sakai, K., Komaki, H., and Gonoi, T. (2015) Identification and Functional Analysis of the Nocardithiocin Gene Cluster in *Nocardia pseudobrasiliensis*. PloS One, 10, e0143264, doi: 10.1371/journal.pone.0143264

42. Harent, S., Vuotto, F., Wallet, F., Flateau, C., Chopin, M. C., Faure, K., and Guery, B. (2013) Pneumonie à *Nocardia pseudobrasiliensis* chez un patient transplanté cardiaque [*Nocardia pseudobrasiliensis* pneumonia in a heart transplant recipient], Med. Mal. Infect., 43, 85–87, doi: 10.1016/j.medmal.2013.01.012

43. Lebeaux, D., Lanternier, F., Degand, N., Catherinot, E., Podglajen, I., Rubio, M. T., Suarez, F., Lecuit, M., Mainardi, J. L., and Lortholary, O. (2010) *Nocardia pseudobrasiliensis* as an emerging cause of opportunistic infection after allogeneic hematopoietic stem cell transplantation, J. Clin. Microbiol., 48, 656–659, doi: 10.1128/JCM.01244-09

44. Veerappan Kandasamy, V., Nagabandi, A., Horowitz, E. A., and Vivekanandan, R. (2015) Multidrug-resistant *Nocardia pseudobrasiliensis* presenting as multiple muscle abscesses, BMJ Case Rep., 2015, bcr2014205262, doi: 10.1136/bcr-2014-205262

45. Seol, C. A., Sung, H., Kim, D. H., Ji, M., Chong, Y. P., and Kim, M. N. (2013) The first Korean case of disseminated mycetoma caused by *Nocardia pseudobrasiliensis* in a patient on long-term corticosteroid therapy for the treatment of microscopic polyangiitis, Ann. Lab. Med., 33, 203–207, doi: 10.3343/alm.2013.33.3.203

46. Zhu, J. W., Zhou, H., Jia, W. Q., You, J., and Xu, R. X. (2020) A clinical case report of brain abscess caused by *Nocardia brasiliensis* in a non-immunocompromised patient and a relevant literature review, BMC Infect. Dis., 20, 328, doi: 10.1186/s12879-020-05052-0

47. Mangieri, N. A., Guevara Nuñez, D., Echavarría, G., Bertona, E., Castello, L., Benchetrit, G., and De Paulis, A. N. (2021) Nocardiosis esporotricoide por *Nocardia brasiliensis* [Sporotrichoid nocardiosis by Nocardia brasiliensis], Rev. Argent. Microbiol., 53, 43–47, doi: 10.1016/j.ram.2020.06.007

48. Verma, P. and Jha, A. (2019) Mycetoma: reviewing a neglected disease, Clin. Exp. Dermatol., 44, 123–129, doi: 10.1111/ced.13642

49. Salinas-Carmona M. C. (2000) *Nocardia brasiliensis*: from microbe to human and experimental infections, Microbes Infect., 2, 1373–1381, doi: 10.1016/s1286-4579(00)01291-0

50. Johansen, M. D., Herrmann, J. L., and Kremer, L. (2020) Non-tuberculous mycobacteria and the rise of *Mycobacterium abscessus*. Nat. Rev. Microbiol., 18, 392–407, doi: 10.1038/s41579-020-0331-1

51. Meir, M., Barkan, D. (2020) Alternative and Experimental Therapies of *Mycobacterium abscessus* Infections, Int. J. Mol. Sci., 21, 6793, doi: 10.3390/ijms21186793

52. Strnad, L., Winthrop, K. L. (2018) Treatment of *Mycobacterium abscessus* Complex, Semin. Respir. Crit. Care Med., 39, 362–376, doi: 10.1055/s-0038-1651494

53. Yonekawa, A., Miyake, N., Minami, J., Murakami, D., Fukano, H., Hoshino, Y., Kubo, K., Chong, Y., Akashi, K., and Shimono, N. (2022) Parotitis caused by *Mycobacteroides abscessus* subspecies *abscessus*, Auris, Nasus, Larynx, 49, 525–528, doi: 10.1016/j.anl.2020.11.005

54. Degiacomi, G., Sammartino, J. C., Chiarelli, L. R., Riabova, O., Makarov, V., and Pasca, M. R. (2019) *Mycobacterium abscessus*, an Emerging and Worrisome Pathogen among Cystic Fibrosis Patients, Int. J. Mol. Sci., 20, 5868, doi: 10.3390/ijms20235868

55. Yoshida, S., Morizumi, S., Sumitomo, K., and Shinohara, T. (2021) Tracheobronchopathia Osteochondroplastica Complicated with *Mycobacteroides abscessus* Pulmonary Disease, Intern. Med., 60, 3051–3052, doi: 10.2169/internalmedicine.6844-20

56. De Carvalho, C. C., da Fonseca, M. M. (2005) The remarkable *Rhodococcus erythropolis*, Appl. Microbiol. Biotechnol., 67, 715–726, doi: 10.1007/s00253-005-1932-3

57. Ma, X., Duan, D., Wang, X., Cao, J., Qiu, J., and Xie, B. (2021) Degradation of *Rhodococcus erythropolis* SY095 modified with functional magnetic Fe_3_O_4_ nanoparticles, R. Soc. Open Sci., 8, 211172, doi: 10.1098/rsos.211172

58. Korzhenkov, A. A., Bakhmutova, E. D., Izotova, A. O., Bavtushnyi, A. A., Sidoruk, K. V., Patrusheva, E. V., Patrushev, M. V., and Toshchakov, S. V. (2021) Draft Genome Sequence of *Rhodococcus erythropolis* VKPM Ac-1659, a Putative Oil-Degrading Strain Isolated from Polluted Soil in Siberia, Microbiol. Resour. Announc., 10, e0053521, doi: 10.1128/MRA.00535-21

59. Thatcher, L. F., Myers, C. A., O’Sullivan, C. A., and Roper, M. M. (2017) Draft Genome Sequence of *Rhodococcus* sp. Strain 66b, Genome Announc., 5, e00229-17, doi: 10.1128/genomeA.00229-17

60. Cunningham-Oakes, E., Pointon, T., Murphy, B., Connor, T. R., and Mahenthiralingam, E. (2020) Genome Sequence of *Pluralibacter gergoviae* ECO77, a Multireplicon Isolate of Industrial Origin, Microbiol. Resour. Announc., 9, e01561–19, doi: 10.1128/MRA.01561-19

61. Chan, K. G., Tee, K. K., Yin, W. F., and Tan, J. Y. (2014) Complete Genome Sequence of *Pluralibacter gergoviae* FB2, an N-Acyl Homoserine Lactone-Degrading Strain Isolated from Packed Fish Paste, Genome Announc., 2, e01276–14, doi: 10.1128/genomeA.01276-14

62. Khashei, R., Edalati Sarvestani, F., Malekzadegan, Y., and Motamedifar, M. (2020) The first report of *Enterobacter gergoviae* carrying *bla*NDM-1 in Iran, Iran. J. Basic Med. Sci., 23, 1184–1190, doi: 10.22038/ijbms.2020.41225.9752

63. Périamé, M., Pagès, J. M., and Davin-Regli, A. (2015) *Enterobacter gergoviae* membrane modifications are involved in the adaptive response to preservatives used in cosmetic industry, J. Appl. Microbiol., 118, 49–61, doi: 10.1111/jam.12676

64. Périamé, M., Philippe, N., Condell, O., Fanning, S., Pagès, J. M., and Davin-Regli, A. (2015) Phenotypic changes contributing to *Enterobacter gergoviae* biocide resistance, Lett. Appl. Microbiol., 61, 121–129, doi: 10.1111/lam.12435

65. Kesieme, E. B., Kesieme, C. N., Akpede, G. O., Okonta, K. E., Dongo, A. E., Gbolagade, A. M., and Eluehike, S. U. (2012) Tension Pneumatocele due to *Enterobacter gergoviae* Pneumonia: A Case Report, Case Rep. Med., 2012, 808630, doi: 10.1155/2012/808630

66. Freire, M. P., De Oliveira Garcia, D., Cury, A. P., Spadão, F., Di Gioia, T. S., Francisco, G. R., Bueno, M. F., Tomaz, M., De Paula, F. J., De Faro, L. B., Piovesan, A. C., Rossi, F., Levin, A. S., David Neto, E., Nahas, W. C., and Pierrotti, L. C. (2016) Outbreak of IMP-producing carbapenem-resistant *Enterobacter gergoviae* among kidney transplant recipients, J. Antimicrob. Chemother., 71, 2577–2585, doi: 10.1093/jac/dkw165

67. Shinjo, R., Uesaka, K., Ihara, K., Loshakova, K., Mizuno, Y., Yano, K., and Tanaka, A. (2016) Complete Genome Sequence of *Kosakonia sacchari* Strain BO-1, an Endophytic Diazotroph Isolated from a Sweet Potato, Genome Announc., 4, e00868–16, doi: 10.1128/genomeA.00868-16

68. Mezzatesta, M. L., Gona, F., and Stefani, S. (2012) *Enterobacter cloacae* complex: clinical impact and emerging antibiotic resistance, Future Microbiol., 7, 887–902, doi: 10.2217/fmb.12.61

69. Ranawat, B., Mishra, S., and Singh, A. (2021) *Enterobacter hormaechei* (MF957335) enhanced yield, disease and salinity tolerance in tomato, Arch. Microbiol., 203, 2659– 2667, doi: 10.1007/s00203-021-02226-5

70. Ranawat, B., Bachani, P., Singh, A., and Mishra, S. (2021). *Enterobacter hormaechei* as Plant Growth-Promoting Bacteria for Improvement in *Lycopersicum esculentum*, Curr. Microbiol., 78, 1208–1217, doi: 10.1007/s00284-021-02368-1

71. Zhang, Q., Wang, S., Zhang, X., Zhang, K., Liu, W., Zhang, R., and Zhang, Z. (2021) *Enterobacter hormaechei* in the intestines of housefly larvae promotes host growth by inhibiting harmful intestinal bacteria, Parasit. Vectors, 14, 598, doi: 10.1186/s13071-021-05053-1

72. Gou, J. J., Liu, N., Guo, L. H., Xu, H., Lv, T., Yu, X., Chen, Y. B., Guo, X. B., Rao, Y. T., and Zheng, B. W. (2020) Carbapenem-Resistant *Enterobacter hormaechei* ST1103 with IMP-26 Carbapenemase and ESBL Gene *bla*_SHV-178_, Infect. Drug Resist., 13, 597–605, doi: 10.2147/IDR.S232514

73. Monowar, T., Rahman, M. S., Bhore, S. J., and Sathasivam, K. V. (2021) Endophytic Bacteria *Enterobacter hormaechei* Fabricated Silver Nanoparticles and Their Antimicrobial Activity, Pharmaceutics, 13, 511, doi: 10.3390/pharmaceutics13040511

74. Wang, Z., Duan, L., Liu, F., Hu, Y., Leng, C., Kan, Y., Yao, L., and Shi, H. (2020) First report of *Enterobacter hormaechei* with respiratory disease in calves, BMC Vet. Res., 16, 1, doi: 10.1186/s12917-019-2207-z

75. Bonnin, R. A., Girlich, D., Jousset, A. B., Emeraud, C., Creton, E., Gauthier, L., Jové, T., Dortet, L., and Naas, T. (2021) Genomic analysis of VIM-2-producing *Enterobacter hormaechei* subsp. *steigerwaltii*, Int. J. Antimicrob. Agents, 57, 106285, doi: 10.1016/j.ijantimicag.2021.106285

76. Gao, W., Howden, B. P., and Stinear, T. P. (2018) Evolution of virulence in *Enterococcus faecium*, a hospital-adapted opportunistic pathogen, Curr. Opin. Microbiol., 41, 76–82, doi: 10.1016/j.mib.2017.11.030

77. Gök, Ş. M., Türk Dağı, H., Kara, F., Arslan, U., and Fındık, D. (2020) Klinik Örneklerden İzole Edilen *Enterococcus faecium* ve *Enterococcus faecalis* İzolatlarının Antibiyotik Direnci ve Virülans Faktörlerinin Araştırılması [Investigation of Antibiotic Resistance and Virulence Factors of *Enterococcus faecium* and *Enterococcus faecalis* Strains Isolated from Clinical Samples], Mikrobiyol. Bul., 54, 26–39, doi: 10.5578/mb.68810

78. Freitas, A. R., Pereira, A. P., Novais, C., and Peixe, L. (2021) Multidrug-resistant high-risk *Enterococcus faecium* clones: can we really define them? Int. J. Antimicrob. Agents, 57, 106227, doi: 10.1016/j.ijantimicag.2020.106227

79. Trościańczyk, A., Nowakiewicz, A., Gnat, S., Łagowski, D., Osińska, M., and Chudzik-Rząd, B. (2021) Comparative study of multidrug-resistant *Enterococcus faecium* obtained from different hosts, J. Med. Microbiol., 70, 10.1099/jmm.0.001340, doi: 10.1099/jmm.0.001340

80. van Hal, S. J., Willems, R., Gouliouris, T., Ballard, S. A., Coque, T. M., Hammerum, A. M., Hegstad, K., Westh, H. T., Howden, B. P., Malhotra-Kumar, S., Werner, G., Yanagihara, K., Earl, A. M., Raven, K. E., Corander, J., Bowden, R., and Enterococcal Group (2021) The global dissemination of hospital clones of *Enterococcus faecium*, Genome Med., 13, 52, doi: 10.1186/s13073-021-00868-0

81. Montironi, I. D., Moliva, M. V., Campra, N. A., Raviolo, J. M., Bagnis, G., Cariddi, L. N., and Reinoso, E. B. (2020) Characterization of an *Enterococcus faecium* strain in a murine mastitis model, J. Appl. Microbiol., 128, 1289–1300, doi: 10.1111/jam.14554

82. Yu, Z., Shi, D., Liu, W., Meng, Y., and Meng, F. (2020) Metabolome responses of *Enterococcus faecium* to acid shock and nitrite stress, Biotechnol. Bioeng., 117, 3559– 3571, doi: 10.1002/bit.27497

83. Duarte, B., Pereira, A. P., Freitas, A. R., Coque, T. M., Hammerum, A. M., Hasman, H., Antunes, P., Peixe, L., and Novais, C. (2019) 2CS-CHX^T^ Operon Signature of Chlorhexidine Tolerance among *Enterococcus faecium* Isolates, Appl. Environ. Microbiol., 85, e01589–19, doi: 10.1128/AEM.01589-19

84. Dündar H. (2016) Bacteriocinogenic Potential of *Enterococcus faecium* Isolated from Wine, Probiotics Antimicrob. Proteins, 8, 150–160, doi: 10.1007/s12602-016-9222-1

85. Salazar, G., Almeida, A., and Gómez, M. (2013) Infección de herida traumática por *Cedecea lapagei*: Comunicación de un caso y revisión de la literatura [*Cedecea lapagei* traumatic wound infection: case report and literature review], Rev. Chilena Infectol., 30, 86–89, doi: 10.4067/S0716-10182013000100015

86. Ramaswamy, V. V., Gummadapu, S., and Suryanarayana, N. (2019) Nosocomial pneumonia and sepsis caused by a rare organism *Cedecea lapagei* in an infant and a review of literature, BMJ Case Rep., 12, e229854, doi: 10.1136/bcr-2019-229854

87. Duperret M. E. (2020) Sinusitis caused by a rare organism, *Cedecea lapagei*, BMJ Case Rep., 13, e235331, doi: 10.1136/bcr-2020-235331

88. Biswal, I., Hussain, N. A., and Grover, R. K. (2015) *Cedecea lapagei* in a patient with malignancy: Report of a rare case, J. Cancer Res. Ther., 11, 646, doi: 10.4103/0973-1482.147736

89. Hai, P. D., Dung, N. M., Tot, N. H., Chinh, N. X., Thuyet, B. T., Hoa, L., Son, P. N., and Thanh, L. V. (2020) First report of pneumonia and septic shock caused by *Cedecea lapagei* in Vietnam, New Microbes New Infect., 36, 100698, doi: 10.1016/j.nmni.2020.100698

90. Ahmad, N., Ali, S. M., and Khan, A. U. (2017) First reported New Delhi metallo-β-lactamase-1-producing *Cedecea lapagei*, Int. J. Antimicrob. Agents, 49, 118–119, doi: 10.1016/j.ijantimicag.2016.10.001

91. Chavez Herrera, V. R., Rosas De Silva, M. F., Orendain Alcaraz, H., Ceja Espiritu, G., Carrazco Peña, K., and Melnikov, V. (2018) Death related to *Cedecea lapagei* in a soft tissue bullae infection: a case report, J. Med. Case Rep., 12, 328, doi: 10.1186/s13256-018-1866-x

92. Dalamaga, M., Karmaniolas, K., Arsenis, G., Pantelaki, M., Daskalopoulou, K., Papadavid, E., and Migdalis, I. (2008) *Cedecea lapagei* bacteremia following cement-related chemical burn injury, Burns, 34, 1205–1207, doi: 10.1016/j.burns.2007.09.001

93. Lopez, L. A., Ibarra, B. S., de la Garza, J. A., Rada, F., Nuñez, A. I., and López, M. G. (2013) First reported case of pneumonia caused by *Cedecea lapagei* in America, Braz. J. Infect. Dis., 17, 626–628, doi: 10.1016/j.bjid.2013.03.003

94. Mathews, S. L., Epps, M. J., Blackburn, R. K., Goshe, M. B., Grunden, A. M., and Dunn, R. R. (2019) Public questions spur the discovery of new bacterial species associated with lignin bioconversion of industrial waste, R. Soc. Open Sci., 6, 180748, doi: 10.1098/rsos.180748

95. Town, J., Audy, P., Boyetchko, S. M., and Dumonceaux, T. J. (2016) High-Quality Draft Genome Sequence of Biocontrol Strain *Pantoea sp*. OXWO6B1, Genome Announc., 4, e00582-16, doi: 10.1128/genomeA.00582-16

96. Coutinho, T. A., Venter, S. N. (2009) *Pantoea ananatis*: an unconventional plant pathogen, Mol. Plant Pathol., 10, 325–335, doi: 10.1111/j.1364-3703.2009.00542.x

97. Xue, Y., Hu, M., Chen, S., Hu, A., Li, S., Han, H., Lu, G., Zeng, L., and Zhou, J. (2021) *Enterobacter asburiae* and *Pantoea ananatis* Causing Rice Bacterial Blight in China, Plant Dis., 105, 2078–2088, doi: 10.1094/PDIS-10-20-2292-RE

98. Asselin, J., Bonasera, J. M., Helmann, T. C., Beer, S. V., and Stodghill, P. V. (2021) Complete Genome Sequence Resources for the Onion Pathogen, Pantoea ananatis OC5a, Phytopathology, 111, 1885–1888, doi: 10.1094/PHYTO-09-20-0416-A

99. Choi, O., Kang, B., Lee, Y., Lee, Y., and Kim, J. (2021) *Pantoea ananatis* carotenoid production confers toxoflavin tolerance and is regulated by Hfq-controlled quorum sensing, MicrobiologyOpen, 10, e1143, doi: 10.1002/mbo3.1143

100. Weller-Stuart, T., De Maayer, P., and Coutinho, T. (2017) *Pantoea ananatis*: genomic insights into a versatile pathogen, Mol. Plant Pathol., 18, 1191–1198, doi: 10.1111/mpp.12517

101. Athanasakopoulou, Z., Sofia, M., Giannakopoulos, A., Papageorgiou, K., Chatzopoulos, D. C., Spyrou, V., Petridou, E., Petinaki, E., and Billinis, C. (2022) ESBL-Producing *Moellerella wisconsensis*-The Contribution of Wild Birds in the Dissemination of a Zoonotic Pathogen, Animals (Basel), 12, 340, doi: 10.3390/ani12030340

102. Hickman-Brenner, F. W., Huntley-Carter, G. P., Saitoh, Y., Steigerwalt, A. G., Farmer, J. J., 3rd, and Brenner, D. J. (1984) *Moellerella wisconsensis*, a new genus and species of Enterobacteriaceae found in human stool specimens, J. Clin. Microbiol., 19, 460–463, doi: 10.1128/jcm.19.4.460-463.1984

103. Chilton, N. B., Dergousoff, S. J., Brzezowska, V., Trost, C. N., and Dunlop, D. R. (2020) American Dog Ticks (*Dermacentor variabilis*) as Biological Indicators of an Association between the Enteric Bacterium *Moellerella wisconsensis* and Striped Skunks *(Mephitis mephitis*) in Southwestern Manitoba, Canada. J. Wildl. Dis., 56, 918–921, doi: 10.7589/2019-09-224

104. Cardentey-Reyes, A., Jacobs, F., Struelens, M. J., and Rodriguez-Villalobos, H. (2009) First case of bacteremia caused by *Moellerella wisconsensis*: case report and a review of the literature, Infection, 37, 544–546, doi: 10.1007/s15010-009-8446-3

105. Casalinuovo, F., Musarella, R. (2009) Isolation of *Moellerella wisconsensis* from the lung of a goat, Vet. Microbiol., 138, 401–402, doi: 10.1016/j.vetmic.2009.03.028

106. Aller, A. I., Castro, C., Medina, M. J., González, M. T., Sevilla, P., Morilla, M. D., Corzo, J. E., and Martín-Mazuelos, E. (2009) Isolation of *Moellerella wisconsensis* from blood culture from a patient with acute cholecystitis, Clin. Microbiol. Infect., 15, 1193– 1194, doi: 10.1111/j.1469-0691.2009.03046.x

107. Hugouvieux-Cotte-Pattat, N., Van Gijsegem, F. (2021) Diversity within the *Dickeya zeae* complex, identification of *Dickeya zeae* and *Dickeya oryzae* members, proposal of the novel species *Dickeya parazeae* sp. nov., Int. J. Syst. Evol. Microbiol., 71, 10.1099/ijsem.0.005059, doi: 10.1099/ijsem.0.005059

108. Huang, N., Pu, X., Zhang, J., Shen, H., Yang, Q., Wang, Z., and Lin, B. (2019) In Vitro Formation of *Dickeya zeae* MS1 Biofilm, Curr. Microbiol., 76, 100–107, doi: 10.1007/s00284-018-1593-y

109. Hu, M., Li, J., Chen, R., Li, W., Feng, L., Shi, L., Xue, Y., Feng, X., Zhang, L., and Zhou, J. (2018) *Dickeya zeae* strains isolated from rice, banana and clivia rot plants show great virulence differentials, BMC Microbiol., 18, 136, doi: 10.1186/s12866-018-1300-y

110. Jiang, S., Zhang, J., Yang, Q., Sun, D., Pu, X., Shen, H., Li, Q., Wang, Z., and Lin, B. (2021) Antimicrobial Activity of Natural Plant Compound Carvacrol Against Soft Rot Disease Agent *Dickeya zeae*, Curr. Microbiol., 78, 3453–3463, doi: 10.1007/s00284-021-02609-3

111. Feng, L., Schaefer, A. L., Hu, M., Chen, R., Greenberg, E. P., and Zhou, J. (2019) Virulence Factor Identification in the Banana Pathogen *Dickeya zeae* MS2, Appl. Environ. Microbiol,, 85, e01611–19, doi: 10.1128/AEM.01611-19

112. Boluk, G., Arizala, D., Dobhal, S., Zhang, J., Hu, J., Alvarez, A. M., and Arif, M. (2021) Genomic and Phenotypic Biology of Novel Strains of *Dickeya zeae* Isolated From Pineapple and Taro in Hawaii: Insights Into Genome Plasticity, Pathogenicity, and Virulence Determinants, Front. Plant Sci., 12, 663851, doi: 10.3389/fpls.2021.663851

113. Liang, Z., Huang, L., He, F., Zhou, X., Shi, Z., Zhou, J., Chen, Y., Lv, M., Chen, Y., and Zhang, L. H. (2019) A Substrate-Activated Efflux Pump, DesABC, Confers Zeamine Resistance to *Dickeya zeae*, mBio, 10, e00713–19, doi: 10.1128/mBio.00713-19

114. Hess, S., Suthaus, A., and Melkonian, M. (2015) *"Candidatus Finniella"* (Rickettsiales, Alphaproteobacteria), Novel Endosymbionts of Viridiraptorid Amoeboflagellates (Cercozoa, Rhizaria), Appl. Environ. Microbiol., 82, 659–670, doi: 10.1128/AEM.02680-15

115. Chung, E. J., Park, J. A., Jeon, C. O., and Chung, Y. R. (2015) *Gynuella sunshinyii* gen. nov., sp. nov., an antifungal rhizobacterium isolated from a halophyte, *Carex scabrifolia* Steud, Int. J. Syst. Evol. Microbiol., 65, 1038–1043, doi: 10.1099/ijs.0.000060

116. Nishijima, M., Adachi, K., Katsuta, A., Shizuri, Y., and Yamasato, K. (2013) *Endozoicomonas numazuensis* sp. nov., a gammaproteobacterium isolated from marine sponges, and emended description of the genus Endozoicomonas Kurahashi and Yokota 2007, Int. J. Syst. Evol. Microbiol., 63, 709–714, doi: 10.1099/ijs.0.042077-0

117. Kieffer, N., Poirel, L., Fournier, C., Haltli, B., Kerr, R., and Nordmann, P. (2019) Characterization of PAN-1, a Carbapenem-Hydrolyzing Class B β-Lactamase From the Environmental Gram-Negative *Pseudobacteriovorax antillogorgiicola*, Front. Microbiol., 10, 1673, doi: 10.3389/fmicb.2019.01673

118. McCauley, E. P., Haltli, B., and Kerr, R. G. (2015) Description of *Pseudobacteriovorax antillogorgiicola* gen. nov., sp. nov., a bacterium isolated from the gorgonian octocoral *Antillogorgia elisabethae*, belonging to the family Pseudobacteriovoracaceae fam. nov., within the order Bdellovibrionales, Int. J. Syst. Evol. Microbiol., 65, 522–530, doi: 10.1099/ijs.0.066266-0

119. Mielko, K. A., Jabłoński, S. J., Milczewska, J., Sands, D., Łukaszewicz, M., and Młynarz, P. (2019) Metabolomic studies of *Pseudomonas aeruginosa*, World J. Microbiol. Biotechnol., 35, 178, doi: 10.1007/s11274-019-2739-1

120. Chevalier, S., Bouffartigues, E., Bodilis, J., Maillot, O., Lesouhaitier, O., Feuilloley, M., Orange, N., Dufour, A., and Cornelis, P. (2017) Structure, function and regulation of *Pseudomonas aeruginosa* porins, FEMS Microbiol. Rev., 41, 698–722, doi: 10.1093/femsre/fux020

121. Sharma, G., Rao, S., Bansal, A., Dang, S., Gupta, S., and Gabrani, R. (2014) *Pseudomonas aeruginosa* biofilm: potential therapeutic targets, Biologicals, 42, 1–7, doi: 10.1016/j.biologicals.2013.11.001

122. Jurado-Martín, I., Sainz-Mejías, M., and McClean, S. (2021) *Pseudomonas aeruginosa*: An Audacious Pathogen with an Adaptable Arsenal of Virulence Factors, Int. J. Mol. Sci., 22, 3128, doi: 10.3390/ijms22063128

123. Kang, D., Kirienko, N. V. (2018) Interdependence between iron acquisition and biofilm formation in *Pseudomonas aeruginosa*, J. Microbiol., 56, 449–457, doi: 10.1007/s12275-018-8114-3

124. Skariyachan, S., Sridhar, V. S., Packirisamy, S., Kumargowda, S. T., and Challapilli, S. B. (2018) Recent perspectives on the molecular basis of biofilm formation by *Pseudomonas aeruginosa* and approaches for treatment and biofilm dispersal, Folia Microbiol. (Praha), 63, 413–432, doi: 10.1007/s12223-018-0585-4

125. Broberg, A., Menkis, A., Vasiliauskas, R. (2006) Kutznerides 1-4, depsipeptides from the actinomycete *Kutzneria sp*. 744 inhabiting mycorrhizal roots of *Picea abies* seedlings, J. Nat. Prod., 69, 97–102, doi: 10.1021/np050378g

126. Jiang, W., Heemstra, J. R., Jr, Forseth, R. R., Neumann, C. S., Manaviazar, S., Schroeder, F. C., Hale, K. J., Walsh, C. T. (2011) Biosynthetic chlorination of the piperazate residue in kutzneride biosynthesis by KthP, Biochemistry, 50, 6063–6072, doi: 10.1021/bi200656k

127. Setser, J. W., Heemstra, J. R., Jr, Walsh, C. T., Drennan, C. L. (2014) Crystallographic evidence of drastic conformational changes in the active site of a flavin-dependent N-hydroxylase, Biochemistry, 53, 6063–6077, doi: 10.1021/bi500655q

128. Duangmal, K., Thamchaipenet, A., Matsumoto, A., Takahashi, Y. (2009) *Pseudonocardia acaciae* sp. nov., isolated from roots of *Acacia auriculiformis* A. Cunn. ex Benth, Int. J. Syst. Evol. Microbiol., 59, 1487–1491, doi: 10.1099/ijs.0.007724-0

129. Labeda, D. P., Donahue, J. M., Sells, S. F., and Kroppenstedt, R. M. (2007) *Lentzea kentuckyensis* sp. nov., of equine origin, Int. J. Syst. Evol. Microbiol., 57, 1780–1783, doi: 10.1099/ijs.0.64245-0

130. Fukami, K., Takagi, F., Shimizu, S., Ishigo, K., Takahashi, M., and Horikawa, T. (2021) Isolation of bacteria able to degrade poly-hydroxybutyrate-co-hydroxyhexanoate, and the inhibitory effects of the degradation products on shrimp pathogen *Vibrio penaeicida*, Microb. Pathog., 160, 105167, doi: 10.1016/j.micpath.2021.105167

131. Fukami, K., Takagi, F., Sonoda, K., Okamoto, H., Kaneno, D., Horikawa, T., Takita, M. (2021) Effects of the Monomeric Components of Poly-hydroxybutyrate-co-hydroxyhexanoate on the Growth of *Vibrio penaeicida* In Vitro and on the Survival of Infected Kuruma Shrimp (*Marsupenaeus japonicus*), Animals (Basel), 11, 567, doi: 10.3390/ani11020567

132. Kawato, S., Nozaki, R., Kondo, H., and Hirono, I. (2018) Draft Genome Sequence of *Vibrio penaeicida* Strain TUMSAT-NU1, Isolated from Diseased Shrimp in Japan, Genome Announc., 6, e00153–18, doi: 10.1128/genomeA.00153-18

133. Goarant, C., Merien, F. (2006) Quantification of *Vibrio penaeicida*, the etiological agent of Syndrome 93 in New Caledonian shrimp, by real-time PCR using SYBR Green I chemistry, J. Microbiol. Methods, 67, 27–35, doi: 10.1016/j.mimet.2006.02.013

134. Aguirre-Guzmán, G., Ascencio, F., and Saulnier, D. (2005) Pathogenicity of *Vibrio penaeicida* for white shrimp *Litopenaeus vannamei*: a cysteine protease-like exotoxin as a virulence factor, Dis. Aquat. Organ., 67, 201–207, doi: 10.3354/dao067201

135. Kadowaki, T., Inagawa, H., Kohchi, C., Nishizawa, T., Takahashi, Y., and Soma, G. (2011) Anti-lipopolysaccharide factor evokes indirect killing of virulent bacteria in kuruma prawn, In Vivo, 25, 741–744.

136. Thompson, F. L., Hoste, B., Thompson, C. C., Goris, J., Gomez-Gil, B., Huys, L., De Vos, P., and Swings, J. (2002) *Enterovibrio norvegicus* gen. nov., sp. nov., isolated from the gut of turbot (*Scophthalmus maximus*) larvae: a new member of the family Vibrionaceae, Int. J. Syst. Evol. Microbiol., 52, 2015–2022, doi: 10.1099/00207713-52-6-2015

137. Thompson, F. L., Thompson, C. C., Naser, S., Hoste, B., Vandemeulebroecke, K., Munn, C., Bourne, D., and Swings, J. (2005) *Photobacterium rosenbergii* sp. nov. and *Enterovibrio coralii* sp. nov., vibrios associated with coral bleaching, Int. J. Syst. Evol. Microbiol., 55, 913–917, doi: 10.1099/ijs.0.63370-0

138. Pascual, J., Macián, M. C., Arahal, D. R., Garay, E., and Pujalte, M. J. (2009) Description of *Enterovibrio nigricans* sp. nov., reclassification of Vibrio calviensis as Enterovibrio calviensis comb. nov. and emended description of the genus Enterovibrio Thompson et al. 2002, Int. J. Syst. Evol. Microbiol, 59, 698–704, doi: 10.1099/ijs.0.001990-0

139. Bastian, M., Heymann, S., and Jacomy, M. (2009) Gephi: an open source software for exploring and manipulating networks, International AAAI Conference on Weblogs and Social Media.

140. Katoh, K., Rozewicki, J., and Yamada, K. D. (2019) MAFFT online service: multiple sequence alignment, interactive sequence choice and visualization, Brief. Bioinform., 20, 1160–1166, doi: 10.1093/bib/bbx108

141. Kumar, S., Stecher, G., Li, M., Knyaz, C., and Tamura, K. (2018) MEGA X: molecular evolutionary genetics analysis across computing platforms, Mol. Biol. Evol., 35, 1547, doi: 10.1093/molbev/msy096

142. Letunic, I. and Bork, P. (2021) Interactive Tree Of Life (iTOL) v5: an online tool for phylogenetic tree display and annotation, Nucleic Acids Res., 49, W293–W296, doi: 10.1093/nar/gkab301

143. Huson, D. H. and Bryant, D. (2006) Application of phylogenetic networks in evolutionary studies, Mol. Biol. Evol., 23, 254–267, doi: 10.1093/molbev/msj030

144. Ashkenazy, H., Abadi, S., Martz, E., Chay, O., Mayrose, I., Pupko, T., and Ben-Tal, N. (2016) ConSurf 2016: an improved methodology to estimate and visualize evolutionary conservation in macromolecules, Nucleic Acids Res., 44, W344–W350, doi: 10.1093/nar/gkw408

145. Berezin, C., Glaser, F., Rosenberg, J. et al. (2004) ConSeq: the identification of functionally and structurally important residues in protein sequences, Bioinformatics, 20, 1322–1324, doi: 10.1093/bioinformatics/bth070

146. Perkins, S. J. (1986) Protein volumes and hydration effects: the calculations of partial specific volumes, neutron scattering matchpoints and 280-nm absorption coefficients for proteins and glycoproteins from amino acid sequences, Eur. J. Biochem., 157, 169–180, doi: 10.1111/j.1432-1033.1986.tb09653.x

147. Hammer, Ø., Harper, D. A., and Ryan, P. D. (2001) PAST: Paleontological statistics software package for education and data analysis, Palaeontologia Electronica, 4, 9.

148. Kim, S. E., Lee, J., An, J. U., Kim, T.-H., Oh, C.-W., Ko, Y.-J., Krishnan, M., Choi, J., Yoon, D.-Y., Kim, Y., and Oh D.-K. (2022) Regioselectivity of an arachidonate 9S-lipoxygenase from *Sphingopyxis macrogoltabida* that biosynthesizes 9S, 15S-and 11S, 17S-dihydroxy fatty acids from C20 and C22 polyunsaturated fatty acids, Biochim. Biophys. Acta Mol. Cell Biol. Lipids, 1867, 159091, doi: 10.1016/j.bbalip.2021.159091

149. Ivanov, I., Heydeck, D., Hofheinz, K., Roffeis, J., O’Donnell, V. B., Kuhn, H., and Walther, M. (2010) Molecular enzymology of lipoxygenases, Arch. Biochem. Biophys., 503, 161–174, doi: 10.1016/j.abb.2010.08.016

150. Morello, E., Pérez-Berezo, T., Boisseau, C., Baranek, T., Guillon, A., Bréa, D., Lanotte, P., Carpena, X., Pietrancosta, N., Hervé, V., Ramphal, R., Cenac, N., and Si-Tahar, M. (2019) *Pseudomonas aeruginosa* lipoxygenase LoxA contributes to lung infection by altering the host immune lipid signaling, Front. Microbiol., 10, 1826, doi: 10.3389/fmicb.2019.01826

151. Banthiya, S., Kalms, J., Yoga, E. G., Ivanov, I., Carpena, X., Hamberg, M., Kuhn, H., Sheerer, P. (2016) Structural and functional basis of phospholipid oxygenase activity of bacterial lipoxygenase from *Pseudomonas aeruginosa*, Biochim. Biophys. Acta Mol. Cell Biol. Lipids, 1861, 1681–1692, doi: 10.1016/j.bbalip.2016.08.002

152. Vidal-Mas, J., Busquets, M., and Manresa, A. (2005) Cloning and expression of a lipoxygenase from *Pseudomonas aeruginosa* 42A2, Antonie van Leeuwenhoek, 87, 245–251, doi: 10.1007/s10482-004-4021-1

153. An, J. U., Kim, B. J., Hong, S. H., and Oh, D. K. (2015) Characterization of an omega-6 linoleate lipoxygenase from *Burkholderia thailandensis* and its application in the production of 13-hydroxyoctadecadienoic acid, Appl. Microbiol. Biotechnol., 99, 5487–5497, doi: 10.1007/s00253-014-6353-8

154. Kim, M. J., Lee, J., Kim, S. E., Shin, K. C., and Oh, D. K. (2023) Production of C20 9S-and C22 11S-hydroxy fatty acids by cells expressing *Shewanella hanedai* arachidonate 9S-lipoxygenase, Appl. Microbiol. Biotechnol., 107, 247–260, doi: 10.1007/s00253-022-12285-3

155. Zhang, B., Chen, M., Xia, B., Lu, Z., Khoo, K. S., Show, P. L., and Lu, F. (2022) Characterization and Preliminary Application of a Novel Lipoxygenase from *Enterovibrio norvegicus*, Foods, 11, 2864, doi: 10.3390/foods11182864

156. Dar, H. H., Tyurina, Y. Y., Mikulska-Ruminska, K., Shrivastava, I., Ting, H. C., Tyurin, V. A., Krieger, J., St Croix, C. M., Watkins, S., Bayir, E., Mao, G., Armbruster, C. R., Kapralov, A., Wang, H., Parsek, M. R., Anthonymuthu, T. S., Ogunsola, A.F., Flitter, B. A., Freedman, C. J., Gaston, J. R., Holman, T. R., Pilewski, J. M., Greenberger, J. S., Mallampalli, R. K., Doi, Y., Lee, J. S., Bahar, I., Bomberger, J. M., Bayır, H., and Kagan, V. E. (2019) *Pseudomonas aeruginosa* utilizes host polyunsaturated phosphatidylethanolamines to trigger theft-ferroptosis in bronchial epithelium, J. Clin. Invest., 128, 4639–4653, doi: 10.1172/JCI99490

157. Aldrovandi, M., Banthiya, S., Meckelmann, S., Zhou, Y., Heydeck, D., O’Donnell, V. B., and Kuhn, H. (2018) Specific oxygenation of plasma membrane phospholipids by *Pseudomonas aeruginosa* lipoxygenase induces structural and functional alterations in mammalian cells, Biochim. Biophys. Acta Mol. Cell Biol. Lipids, 1863, 152–164, doi: 10.1016/j.bbalip.2017.11.005

158. Beccaccioli, M., Pucci, N., Salustri, M., Scortichini, M., Zaccaria, M., Momeni, B., Loreti, S., Reverberi, M., and Scala, V. (2022) Fungal and bacterial oxylipins are signals for intra-and inter-cellular communication within plant disease, Front. Plant Sci., 13, 823233, doi: 10.3389/fpls.2022.823233

159. Wang, K. D., Borrego, E. J., Kenerley, C. M., and Kolomiets, M. V. (2020) Oxylipins other than jasmonic acid are xylem-resident signals regulating systemic resistance induced by *Trichoderma virens* in maize, Plant Cell, 32, 166–185, doi: 10.1105/tpc.19.00487

160. Dawkins, R. (1976) The Selfish Gene, New York: Oxford University Press.

161. Yan, C. and Xie, D. (2015) Jasmonate in plant defence: sentinel or double agent? Plant Biotechnol. J., 13, 1233–1240, doi: 10.1111/pbi.12417

162. Chini, A., Cimmino, A., Masi, M., Reveglia, P., Nocera, P., Solano, R., and Evidente, A. (2018) The fungal phytotoxin lasiojasmonate A activates the plant jasmonic acid pathway, J. Exp. Bot., 69, 3095–3102, doi: 10.1093/jxb/ery114

163. Zheng, D., Liwinski, T., and Elinav, E. (2020) Interaction between microbiota and immunity in health and disease, Cell Res., 30, 492–506, doi: 10.1038/s41422-020-0332-7

164. Kurakin, G. (2021) Bacterial oxylipins: a key to multicellularity and to combating antimicrobial resistance? Springer Nature Research Communities, URL: https://communities.springernature.com/posts/bacterial-oxylipins-a-key-to-multicellularity-and-to-combating-antimicrobial-resistance, accessed on 24.02.2024.

