## Supplementary Tables for "Bacterial lipoxygenases are associated with host-microbe interactions and may provide cross-kingdom host jumps"

**Table S1.** Bacterial species for the network text analysis along with corresponding accession IDs of lipoxygenases (if multiple sequences are present in the database, only one is specified) and PubMed IDs (PMIDs) of articles included in the network text analysis (if none given, the justification if briefly specified). The "Phylogenetics" column specifies whether this species was included in the phylogenetic analysis or not.

| Species | Source | Accession ID | Phylogenetics | PMIDs of the species |
|---|---|---|---|---|
| *Dickeya zeae* | NCBI | WP_033581576.1 | No | 34726587, 30390102, 30336787, 34263355, 31540986, 34456933, 31138747 |
| *Rhodococcus sp. 66b* | NCBI | WP_019749966.1 | No | 28546474 |
| *Gynuella sunshinyii* | NCBI | WP_044618294.1 | No | 25575829 |
| *Pseudonocardia acaciae* | NCBI | WP_169747922.1 | No | 19502340 |
| *Enterobacter hormaechei* | NCBI | KAA0874857.1 | No | 22827309, 33712862, 33625569, 34876203, 32110070, 33917798, 31900161, 33493673 |
| *Candidatus Finniella inopinata* | NCBI | WP_130153935.1 | No | 26567303 |
| *Candidatus Entotheonella palauensis* | NCBI | WP_089937704.1 | No | 33038718 (No appropriate ecological data). |
| *Enterovibrio calviensis* | UniProt | A0A1E5BLH8_9GAMM | No | 19329591 |
| *Enterovibrio coralii* | UniProt | A0A135I5H1_9GAMM | No | 15774685 |
| *Enterovibrio nigricans* | UniProt | A0A1T4W935_9GAMM | No | 19329591 (This species was not analyzed since it shared the only PubMed article with already analyzed *E. calviensis*). |

| | | | | |
|---|---|---|---|---|
| *Cedecea lapagei* | UniProt | A0A447V3N2_9ENTR | No | 23450417, 31300600, 32690571, 26458603, 32612841, 28244375, 30388965, 18467034, 24035464, 31031986 |
| *Enterococcus faecium* | NCBI | PWS22692.1 | No | 29227922, 32050876, 33207280, 33207280, 33750516, 33785076, 31840319, 32662876, 31562170, 27406790 |
| *Burkholderia stagnalis* | UniProt | A0A3P0FWH9_9BURK | No | No (No appropriate ecological data). |
| *Pseudomonas aeruginosa* | UniProt | LOXA_PSEAE | Yes | 31701321, 28981745, 24309094, 33803907, 29948830, 29352409 |
| *Burkholderia singularis* | UniProt | A0A238H4J7_9BURK | Yes | 28932212 |
| *Burkholderia thailandensis* | UniProt | Q2SW25_BURTA | Yes | 32281050, 33008823, 28510362, 32900830, 32150572, 23209596, 27517336 |
| *Burkholderia gladioli* | UniProt | A0A2S4NUU5_BURGA | Yes | 33174713, 33297590, 32118858, 30342825, 22648018, 33459584, 29509962, 12131039, 16148855, 25926303 |
| *Variovorax guangxiensis* | UniProt | A0A433MUY6_9BURK | Yes | 25504533 |
| *Variovorax gossypii* | UniProt | A0A3S0GW99_9BURK | Yes | 26341669 |

| | | | | |
|---|---|---|---|---|
| *Variovorax paradoxus* | UniProt | A0A0P8ZCL8_VARPD | Yes | 33433645, 31899322 |
| *Nocardia seriolae* | UniProt | A0A1I9ZED9_9NOCA | Yes | 33150871, 30117297, 30117618, 19335609, 32959416, 32196698, 28257489 |
| *Nocardia brasiliensis* | UniProt | A0A6G9XMP6_9NOCA | Yes | 32381049, 32739070, 29808607, 11018454 |
| *Nocardia pseudobrasiliensis* | UniProt | A0A370IBC4_9NOCA | Yes | 32602372, 8573505, 26588225, 23465630, 19940053, 25948839, 23667849 |
| *Kosakonia sp. AG348* | UniProt | A0A369A5J2_9ENTR | Yes | No (Zero PubMed search results). |
| *Kosakonia sacchari* | UniProt | A0A6P1QMY5_9ENTR | Yes | 27609910 |
| *Pantoea ananas* | UniProt | A0A7H8I474_PANAN | Yes | 19400836, 33342235, 33724871, 33269542, 27880983 |
| *Pantoea sp. OXWO6B1* | NCBI | WP_063877190.1 | Yes | 27340064 |
| *Pluralibacter gergoviae* | UniProt | A0A251ZKI4_PLUGE | Yes | 32107303, 25502672, 32963740, 25355161, 25919899, 23056055, 27197663 |
| *Moellerella wisconsensis* | NCBI | WP_052956414.1 | Yes | 35158664, 6715516, 32402233, 19730786, 19394768, 19732083 |
| *Enterovibrio norvegicus* | NCBI | WP_017014538.1 | Yes | 12508862 |
| *Lentzea kentuckyensis* | NCBI | WP_158102866.1 | Yes | 17684256 |
| *Kutzneria sp. 744* | NCBI | WP_052395874.1 | Yes | 16441076, 21648411, 25184411 |

| | | | | |
|---|---|---|---|---|
| *Mycobacteroides abscessus* | NCBI | WP_074347916.1 | Yes | 32086501, 32948001, 30071551, 33246745, 31766758, 33814496 |
| *Pseudobacteriovorax antillogorgiicola* | NCBI | WP_132324275.1 | Yes | 31396187, 25389148 |
| *Endozoicomonas numazuensis* | NCBI | WP_034835369.1 | No | 22544802 |
| *Rhodococcus erythropolis* | NCBI | WP_256065274.1 | No | 15711940, 34950489, 34292073 |

**Table S2.** Full table of terms included in the network text analysis as nodes (along with their corresponding weights).

| id | Weight | Color |
|---|---|---|
| AMR | 9 | yellow |
| human | 8 | red |
| plant | 7 | green |
| emerging | 7 | yellow |
| MDR | 6 | yellow |
| root | 5 | green |
| nosocomial | 5 | red |
| pneumonia | 5 | red |
| lung | 5 | red |
| clinical | 4 | red |
| soil | 5 | gray |
| gut | 5 | gray |
| growth-promoting | 4 | green |
| rhizosphere | 4 | green |
| endophytic | 4 | green |
| healthy | 4 | gray |
| antifungal | 4 | gray |
| bacteremia | 4 | red |
| cultured | 4 | blue |
| opportunistic | 4 | red |
| banana | 3 | green |
| immunocompetent | 3 | red |
| biocontrol | 3 | green |
| cystic fibrosis | 3 | red |
| sepsis | 2 | red |
| animal | 3 | gray |
| insect | 3 | magenta |
| abscess | 3 | red |
| nocardiosis | 3 | gray |
| skin | 3 | red |

| | | |
|---|---|---|
| serious | 2 | yellow |
| aquaculture | 2 | blue |
| chronic | 3 | gray |
| transplant | 3 | red |
| water | 2 | blue |
| biofilm | 3 | gray |
| immunocompromised | 2 | red |
| leaves | 2 | green |
| bloodstream | 2 | red |
| blood | 2 | red |
| invasive | 2 | gray |
| leukemia | 2 | red |
| beneficial | 2 | gray |
| child | 2 | red |
| pulmonary | 2 | red |
| antimicrobial | 2 | gray |
| saprophyte | 2 | gray |
| marine | 2 | blue |
| fish | 2 | blue |
| kidney | 2 | red |
| granulomatous | 2 | red |
| threat | 2 | yellow |
| epizootic | 2 | gray |
| carbapenem-resistant | 2 | yellow |
| mycetoma | 2 | red |
| uncommon | 2 | gray |
| environment | 2 | gray |
| respiratory | 2 | red |
| bioconversion | 2 | gray |
| degradation | 2 | gray |
| industrial | 2 | gray |
| biocide resistance | 2 | yellow |
| urinary | 2 | red |
| larvae | 2 | gray |
| concern | 2 | yellow |
| animals | 2 | gray |
| healthcare | 2 | red |
| rare | 2 | gray |
| compromised | 2 | red |
| NDM-1 | 2 | yellow |
| crops | 2 | green |
| potato | 2 | green |
| rice | 2 | green |
| rot | 2 | green |
| carbapenemase | 2 | yellow |

**Table S3.** Full table of term co-occurrences included in the network text analysis as edges (along with the co-occurrence numbers as weights).

| **Source** | **Target** | **Weight** |
|---|---|---|
| plant | growth-promoting | 1 |
| plant | rhizosphere | 1 |
| plant | endophytic | 1 |
| growth-promoting | rhizosphere | 1 |
| growth-promoting | endophytic | 2 |
| rhizosphere | endophytic | 1 |
| rhizosphere | banana | 1 |
| rhizosphere | plant | 1 |
| banana | plant | 2 |
| root | healthy | 1 |
| nosocomial | immunocompromised | 2 |
| nosocomial | immunocompetent | 1 |
| nosocomial | endophytic | 2 |
| nosocomial | leaves | 1 |
| nosocomial | antifungal | 1 |
| nosocomial | biocontrol | 1 |
| nosocomial | cystic fibrosis | 2 |
| nosocomial | sepsis | 2 |
| nosocomial | bacteremia | 3 |
| nosocomial | immunosuppression | 1 |
| nosocomial | bloodstream | 2 |
| nosocomial | blood | 1 |
| nosocomial | invasive | 1 |
| nosocomial | AMR | 4 |
| nosocomial | leukemia | 2 |
| nosocomial | pneumonia | 3 |
| nosocomial | human | 3 |
| nosocomial | animal | 3 |
| nosocomial | plant | 2 |
| nosocomial | beneficial | 2 |
| nosocomial | lung | 3 |
| nosocomial | insect | 3 |
| nosocomial | child | 1 |
| nosocomial | abscess | 1 |
| nosocomial | pulmonary | 1 |
| immunocompromised | immunocompetent | 1 |
| immunocompromised | endophytic | 1 |
| immunocompromised | leaves | 1 |
| immunocompromised | antifungal | 1 |
| immunocompromised | biocontrol | 1 |
| immunocompromised | cystic fibrosis | 2 |
| immunocompromised | sepsis | 1 |
| immunocompromised | bacteremia | 2 |

| | | |
|---|---|---|
| immunocompromised | immunosuppression | 1 |
| immunocompromised | bloodstream | 1 |
| immunocompromised | blood | 1 |
| immunocompromised | invasive | 1 |
| immunocompromised | AMR | 2 |
| immunocompromised | leukemia | 1 |
| immunocompromised | pneumonia | 2 |
| immunocompromised | human | 2 |
| immunocompromised | animal | 2 |
| immunocompromised | plant | 1 |
| immunocompromised | beneficial | 1 |
| immunocompromised | lung | 2 |
| immunocompromised | insect | 1 |
| immunocompromised | child | 1 |
| immunocompromised | abscess | 1 |
| immunocompromised | pulmonary | 1 |
| immunocompetent | endophytic | 1 |
| immunocompetent | leaves | 1 |
| immunocompetent | antifungal | 1 |
| immunocompetent | biocontrol | 1 |
| immunocompetent | cystic fibrosis | 1 |
| immunocompetent | sepsis | 1 |
| immunocompetent | bacteremia | 1 |
| immunocompetent | immunosuppression | 1 |
| immunocompetent | bloodstream | 1 |
| immunocompetent | blood | 1 |
| immunocompetent | invasive | 2 |
| immunocompetent | AMR | 2 |
| immunocompetent | leukemia | 1 |
| immunocompetent | pneumonia | 2 |
| immunocompetent | human | 2 |
| immunocompetent | animal | 1 |
| immunocompetent | plant | 1 |
| immunocompetent | beneficial | 1 |
| immunocompetent | lung | 1 |
| immunocompetent | insect | 1 |
| immunocompetent | child | 1 |
| immunocompetent | abscess | 3 |
| immunocompetent | pulmonary | 1 |
| endophytic | leaves | 1 |
| endophytic | antifungal | 1 |
| endophytic | biocontrol | 1 |
| endophytic | cystic fibrosis | 1 |
| endophytic | sepsis | 1 |
| endophytic | bacteremia | 1 |
| endophytic | immunosuppression | 1 |

| | | |
|---|---|---|
| endophytic | bloodstream | 2 |
| endophytic | blood | 1 |
| endophytic | invasive | 1 |
| endophytic | AMR | 2 |
| endophytic | leukemia | 1 |
| endophytic | pneumonia | 1 |
| endophytic | human | 1 |
| endophytic | animal | 2 |
| endophytic | plant | 3 |
| endophytic | beneficial | 2 |
| endophytic | lung | 2 |
| endophytic | insect | 2 |
| endophytic | child | 1 |
| endophytic | abscess | 1 |
| endophytic | pulmonary | 1 |
| leaves | antifungal | 2 |
| leaves | biocontrol | 2 |
| leaves | cystic fibrosis | 1 |
| leaves | sepsis | 1 |
| leaves | bacteremia | 1 |
| leaves | immunosuppression | 1 |
| leaves | bloodstream | 1 |
| leaves | blood | 1 |
| leaves | invasive | 1 |
| leaves | AMR | 1 |
| leaves | leukemia | 1 |
| leaves | pneumonia | 1 |
| leaves | human | 2 |
| leaves | animal | 1 |
| leaves | plant | 2 |
| leaves | beneficial | 1 |
| leaves | lung | 1 |
| leaves | insect | 1 |
| leaves | child | 1 |
| leaves | abscess | 1 |
| leaves | pulmonary | 1 |
| antifungal | biocontrol | 2 |
| antifungal | cystic fibrosis | 1 |
| antifungal | sepsis | 1 |
| antifungal | bacteremia | 1 |
| antifungal | immunosuppression | 1 |
| antifungal | bloodstream | 1 |
| antifungal | blood | 1 |
| antifungal | invasive | 1 |
| antifungal | AMR | 1 |
| antifungal | leukemia | 1 |

| | | |
|---|---|---|
| antifungal | pneumonia | 1 |
| antifungal | human | 2 |
| antifungal | animal | 1 |
| antifungal | plant | 2 |
| antifungal | beneficial | 1 |
| antifungal | lung | 1 |
| antifungal | insect | 1 |
| antifungal | child | 1 |
| antifungal | abscess | 1 |
| antifungal | pulmonary | 1 |
| biocontrol | cystic fibrosis | 1 |
| biocontrol | sepsis | 1 |
| biocontrol | bacteremia | 1 |
| biocontrol | immunosuppression | 1 |
| biocontrol | bloodstream | 1 |
| biocontrol | blood | 1 |
| biocontrol | invasive | 1 |
| biocontrol | AMR | 1 |
| biocontrol | leukemia | 1 |
| biocontrol | pneumonia | 1 |
| biocontrol | human | 2 |
| biocontrol | animal | 1 |
| biocontrol | plant | 2 |
| biocontrol | beneficial | 1 |
| biocontrol | lung | 1 |
| biocontrol | insect | 1 |
| biocontrol | child | 1 |
| biocontrol | abscess | 1 |
| biocontrol | pulmonary | 1 |
| cystic fibrosis | sepsis | 1 |
| cystic fibrosis | bacteremia | 2 |
| cystic fibrosis | immunosuppression | 1 |
| cystic fibrosis | bloodstream | 1 |
| cystic fibrosis | blood | 1 |
| cystic fibrosis | invasive | 1 |
| cystic fibrosis | AMR | 3 |
| cystic fibrosis | leukemia | 1 |
| cystic fibrosis | pneumonia | 2 |
| cystic fibrosis | human | 3 |
| cystic fibrosis | animal | 2 |
| cystic fibrosis | plant | 1 |
| cystic fibrosis | beneficial | 1 |
| cystic fibrosis | lung | 3 |
| cystic fibrosis | insect | 1 |
| cystic fibrosis | child | 1 |
| cystic fibrosis | abscess | 1 |

| | | |
|---|---|---|
| cystic fibrosis | pulmonary | 2 |
| sepsis | bacteremia | 2 |
| sepsis | immunosuppression | 1 |
| sepsis | bloodstream | 1 |
| sepsis | blood | 1 |
| sepsis | invasive | 1 |
| sepsis | AMR | 1 |
| sepsis | leukemia | 2 |
| sepsis | pneumonia | 2 |
| sepsis | animal | 1 |
| sepsis | plant | 1 |
| sepsis | beneficial | 1 |
| sepsis | lung | 1 |
| sepsis | insect | 2 |
| sepsis | child | 1 |
| sepsis | abscess | 1 |
| sepsis | pulmonary | 1 |
| bacteremia | immunosuppression | 1 |
| bacteremia | bloodstream | 1 |
| bacteremia | blood | 2 |
| bacteremia | invasive | 1 |
| bacteremia | AMR | 3 |
| bacteremia | leukemia | 2 |
| bacteremia | pneumonia | 3 |
| bacteremia | human | 3 |
| bacteremia | animal | 2 |
| bacteremia | plant | 1 |
| bacteremia | beneficial | 1 |
| bacteremia | lung | 3 |
| bacteremia | insect | 2 |
| bacteremia | child | 1 |
| bacteremia | abscess | 1 |
| bacteremia | pulmonary | 1 |
| immunosuppression | bloodstream | 1 |
| immunosuppression | blood | 1 |
| immunosuppression | invasive | 1 |
| immunosuppression | AMR | 1 |
| immunosuppression | leukemia | 1 |
| immunosuppression | pneumonia | 1 |
| immunosuppression | human | 1 |
| immunosuppression | animal | 1 |
| immunosuppression | plant | 1 |
| immunosuppression | beneficial | 1 |
| immunosuppression | lung | 1 |
| immunosuppression | insect | 1 |
| immunosuppression | child | 1 |

| | | |
|---|---|---|
| immunosuppression | abscess | 1 |
| immunosuppression | pulmonary | 1 |
| bloodstream | blood | 1 |
| bloodstream | invasive | 1 |
| bloodstream | AMR | 2 |
| bloodstream | leukemia | 1 |
| bloodstream | pneumonia | 1 |
| bloodstream | human | 1 |
| bloodstream | animal | 2 |
| bloodstream | plant | 2 |
| bloodstream | beneficial | 2 |
| bloodstream | lung | 2 |
| bloodstream | insect | 2 |
| bloodstream | child | 1 |
| bloodstream | abscess | 1 |
| bloodstream | pulmonary | 1 |
| blood | invasive | 1 |
| blood | AMR | 2 |
| blood | leukemia | 1 |
| blood | pneumonia | 1 |
| blood | human | 2 |
| blood | animal | 1 |
| blood | plant | 1 |
| blood | beneficial | 1 |
| blood | lung | 2 |
| blood | insect | 1 |
| blood | child | 1 |
| blood | abscess | 1 |
| blood | pulmonary | 1 |
| invasive | AMR | 2 |
| invasive | leukemia | 1 |
| invasive | pneumonia | 2 |
| invasive | human | 1 |
| invasive | animal | 1 |
| invasive | plant | 1 |
| invasive | beneficial | 1 |
| invasive | lung | 1 |
| invasive | insect | 1 |
| invasive | child | 1 |
| invasive | abscess | 2 |
| invasive | pulmonary | 1 |
| AMR | leukemia | 1 |
| AMR | pneumonia | 4 |
| AMR | human | 5 |
| AMR | animal | 3 |
| AMR | plant | 2 |

| | | |
|---|---|---|
| AMR | beneficial | 2 |
| AMR | lung | 5 |
| AMR | insect | 2 |
| AMR | child | 2 |
| AMR | abscess | 2 |
| AMR | pulmonary | 2 |
| leukemia | pneumonia | 2 |
| leukemia | human | 1 |
| leukemia | animal | 1 |
| leukemia | plant | 1 |
| leukemia | beneficial | 1 |
| leukemia | lung | 1 |
| leukemia | insect | 2 |
| leukemia | child | 1 |
| leukemia | abscess | 1 |
| leukemia | pulmonary | 1 |
| pneumonia | human | 2 |
| pneumonia | animal | 2 |
| pneumonia | plant | 1 |
| pneumonia | beneficial | 1 |
| pneumonia | lung | 2 |
| pneumonia | insect | 2 |
| pneumonia | child | 2 |
| pneumonia | abscess | 2 |
| pneumonia | pulmonary | 1 |
| human | animal | 2 |
| human | plant | 2 |
| human | beneficial | 1 |
| human | lung | 4 |
| human | insect | 1 |
| human | child | 1 |
| human | abscess | 2 |
| human | pulmonary | 2 |
| animal | plant | 2 |
| animal | beneficial | 2 |
| animal | lung | 3 |
| animal | insect | 2 |
| animal | child | 1 |
| animal | abscess | 1 |
| animal | pulmonary | 1 |
| plant | beneficial | 2 |
| plant | lung | 2 |
| plant | insect | 2 |
| plant | child | 1 |
| plant | abscess | 1 |
| plant | pulmonary | 1 |

| | | |
|---|---|---|
| beneficial | lung | 2 |
| beneficial | insect | 2 |
| beneficial | child | 1 |
| beneficial | abscess | 1 |
| beneficial | pulmonary | 1 |
| lung | insect | 2 |
| lung | child | 1 |
| lung | abscess | 1 |
| lung | pulmonary | 2 |
| insect | child | 1 |
| insect | abscess | 1 |
| insect | pulmonary | 1 |
| child | abscess | 1 |
| child | pulmonary | 1 |
| abscess | pulmonary | 1 |
| human | clinical | 2 |
| soil | antimicrobial | 2 |
| soil | saprophyte | 2 |
| antimicrobial | saprophyte | 1 |
| nocardiosis | marine | 1 |
| nocardiosis | fish | 1 |
| nocardiosis | skin | 2 |
| nocardiosis | kidney | 1 |
| nocardiosis | granulomatous | 1 |
| nocardiosis | serious | 1 |
| nocardiosis | threat | 1 |
| nocardiosis | cultured | 1 |
| nocardiosis | epizootic | 1 |
| nocardiosis | aquaculture | 1 |
| nocardiosis | chronic | 1 |
| nocardiosis | AMR | 2 |
| marine | fish | 1 |
| marine | skin | 1 |
| marine | kidney | 1 |
| marine | granulomatous | 1 |
| marine | serious | 1 |
| marine | threat | 1 |
| marine | cultured | 1 |
| marine | epizootic | 1 |
| marine | aquaculture | 1 |
| marine | chronic | 1 |
| marine | AMR | 1 |
| fish | skin | 1 |
| fish | kidney | 1 |
| fish | granulomatous | 1 |
| fish | serious | 1 |

| | | |
|---|---|---|
| fish | threat | 1 |
| fish | cultured | 2 |
| fish | epizootic | 1 |
| fish | aquaculture | 1 |
| fish | chronic | 1 |
| fish | AMR | 1 |
| skin | kidney | 1 |
| skin | granulomatous | 2 |
| skin | serious | 2 |
| skin | threat | 1 |
| skin | cultured | 1 |
| skin | epizootic | 1 |
| skin | aquaculture | 1 |
| skin | chronic | 2 |
| skin | AMR | 2 |
| kidney | granulomatous | 1 |
| kidney | serious | 1 |
| kidney | threat | 1 |
| kidney | cultured | 1 |
| kidney | epizootic | 1 |
| kidney | aquaculture | 1 |
| kidney | chronic | 1 |
| kidney | AMR | 2 |
| granulomatous | serious | 2 |
| granulomatous | threat | 1 |
| granulomatous | cultured | 1 |
| granulomatous | epizootic | 1 |
| granulomatous | aquaculture | 1 |
| granulomatous | chronic | 2 |
| granulomatous | AMR | 2 |
| serious | threat | 1 |
| serious | cultured | 1 |
| serious | epizootic | 1 |
| serious | chronic | 2 |
| serious | AMR | 2 |
| threat | cultured | 1 |
| threat | epizootic | 1 |
| threat | aquaculture | 1 |
| threat | chronic | 2 |
| threat | AMR | 2 |
| cultured | epizootic | 2 |
| cultured | aquaculture | 2 |
| cultured | chronic | 1 |
| cultured | AMR | 1 |
| epizootic | aquaculture | 2 |
| epizootic | chronic | 1 |

| | | |
|---|---|---|
| epizootic | AMR | 1 |
| aquaculture | chronic | 1 |
| aquaculture | AMR | 1 |
| chronic | AMR | 3 |
| nocardiosis | emerging | 1 |
| nocardiosis | invasive | 1 |
| nocardiosis | immunocompetent | 2 |
| nocardiosis | abscess | 2 |
| nocardiosis | opportunistic | 1 |
| nocardiosis | pneumonia | 1 |
| nocardiosis | transplant | 1 |
| nocardiosis | carbapenem-resistant | 1 |
| nocardiosis | MDR | 1 |
| nocardiosis | mycetoma | 2 |
| emerging | invasive | 1 |
| emerging | immunocompetent | 1 |
| emerging | abscess | 1 |
| emerging | opportunistic | 3 |
| emerging | pneumonia | 2 |
| emerging | transplant | 2 |
| emerging | carbapenem-resistant | 2 |
| emerging | AMR | 4 |
| emerging | MDR | 3 |
| emerging | mycetoma | 1 |
| invasive | opportunistic | 1 |
| invasive | transplant | 1 |
| invasive | carbapenem-resistant | 1 |
| invasive | MDR | 1 |
| invasive | mycetoma | 1 |
| immunocompetent | opportunistic | 1 |
| immunocompetent | transplant | 1 |
| immunocompetent | carbapenem-resistant | 1 |
| immunocompetent | MDR | 1 |
| immunocompetent | mycetoma | 2 |
| abscess | opportunistic | 1 |
| abscess | transplant | 1 |
| abscess | carbapenem-resistant | 1 |
| abscess | MDR | 1 |
| abscess | mycetoma | 2 |
| opportunistic | pneumonia | 1 |
| opportunistic | transplant | 2 |
| opportunistic | carbapenem-resistant | 1 |
| opportunistic | AMR | 4 |
| opportunistic | MDR | 3 |
| opportunistic | mycetoma | 1 |
| pneumonia | transplant | 2 |

| | | |
|---|---|---|
| pneumonia | carbapenem-resistant | 1 |
| pneumonia | MDR | 3 |
| pneumonia | mycetoma | 1 |
| transplant | carbapenem-resistant | 1 |
| transplant | AMR | 3 |
| transplant | MDR | 3 |
| transplant | mycetoma | 1 |
| carbapenem-resistant | AMR | 2 |
| carbapenem-resistant | MDR | 1 |
| carbapenem-resistant | mycetoma | 1 |
| AMR | MDR | 6 |
| AMR | mycetoma | 1 |
| MDR | mycetoma | 1 |
| abscess | soil | 1 |
| abscess | skin | 1 |
| abscess | saprophyte | 1 |
| abscess | uncommon | 1 |
| soil | nocardiosis | 1 |
| soil | immunocompetent | 1 |
| soil | skin | 1 |
| soil | mycetoma | 1 |
| soil | human | 1 |
| soil | uncommon | 1 |
| nocardiosis | saprophyte | 1 |
| nocardiosis | human | 1 |
| nocardiosis | uncommon | 1 |
| immunocompetent | skin | 1 |
| immunocompetent | saprophyte | 1 |
| immunocompetent | uncommon | 1 |
| skin | mycetoma | 1 |
| skin | saprophyte | 1 |
| skin | human | 2 |
| skin | uncommon | 1 |
| mycetoma | saprophyte | 1 |
| mycetoma | human | 1 |
| mycetoma | uncommon | 1 |
| saprophyte | human | 1 |
| saprophyte | uncommon | 1 |
| human | uncommon | 1 |
| AMR | environment | 2 |
| AMR | respiratory | 2 |
| environment | serious | 1 |
| environment | opportunistic | 2 |
| environment | human | 2 |
| environment | respiratory | 1 |
| environment | skin | 1 |

| | | |
|---|---|---|
| environment | chronic | 1 |
| environment | MDR | 2 |
| environment | cystic fibrosis | 1 |
| environment | lung | 1 |
| environment | transplant | 1 |
| environment | pulmonary | 1 |
| environment | granulomatous | 1 |
| environment | emerging | 2 |
| serious | opportunistic | 2 |
| serious | human | 1 |
| serious | respiratory | 1 |
| serious | MDR | 1 |
| serious | cystic fibrosis | 1 |
| serious | lung | 1 |
| serious | transplant | 1 |
| serious | pulmonary | 1 |
| serious | emerging | 1 |
| opportunistic | human | 3 |
| opportunistic | respiratory | 1 |
| opportunistic | skin | 1 |
| opportunistic | chronic | 1 |
| opportunistic | cystic fibrosis | 1 |
| opportunistic | lung | 1 |
| opportunistic | pulmonary | 1 |
| opportunistic | granulomatous | 1 |
| human | respiratory | 1 |
| human | chronic | 2 |
| human | MDR | 4 |
| human | transplant | 1 |
| human | granulomatous | 1 |
| human | emerging | 3 |
| respiratory | skin | 1 |
| respiratory | chronic | 1 |
| respiratory | MDR | 1 |
| respiratory | cystic fibrosis | 1 |
| respiratory | lung | 2 |
| respiratory | transplant | 1 |
| respiratory | pulmonary | 1 |
| respiratory | granulomatous | 1 |
| respiratory | emerging | 2 |
| skin | MDR | 1 |
| skin | cystic fibrosis | 1 |
| skin | lung | 1 |
| skin | transplant | 1 |
| skin | pulmonary | 1 |
| skin | emerging | 1 |

| | | |
|---|---|---|
| chronic | MDR | 2 |
| chronic | cystic fibrosis | 2 |
| chronic | lung | 2 |
| chronic | transplant | 1 |
| chronic | pulmonary | 1 |
| chronic | emerging | 1 |
| MDR | cystic fibrosis | 2 |
| MDR | lung | 3 |
| MDR | pulmonary | 1 |
| MDR | granulomatous | 1 |
| cystic fibrosis | transplant | 1 |
| cystic fibrosis | granulomatous | 1 |
| cystic fibrosis | emerging | 1 |
| lung | transplant | 1 |
| lung | granulomatous | 1 |
| lung | emerging | 2 |
| transplant | pulmonary | 1 |
| transplant | granulomatous | 1 |
| pulmonary | granulomatous | 1 |
| pulmonary | emerging | 1 |
| granulomatous | emerging | 1 |
| bioconversion | degradation | 1 |
| bioconversion | soil | 1 |
| degradation | soil | 2 |
| soil | plant | 1 |
| soil | growth-promoting | 1 |
| plant | growth-promoting | 3 |
| plant | degradation | 1 |
| growth-promoting | degradation | 1 |
| AMR | industrial | 1 |
| AMR | clinical | 3 |
| AMR | biocide resistance | 2 |
| AMR | urinary | 2 |
| industrial | clinical | 1 |
| industrial | biocide resistance | 1 |
| industrial | pneumonia | 2 |
| industrial | child | 1 |
| industrial | kidney | 1 |
| industrial | transplant | 1 |
| industrial | urinary | 1 |
| industrial | MDR | 1 |
| clinical | biocide resistance | 1 |
| clinical | pneumonia | 1 |
| clinical | child | 1 |
| clinical | kidney | 1 |
| clinical | transplant | 1 |

| | | |
|---|---|---|
| clinical | urinary | 1 |
| clinical | MDR | 2 |
| biocide resistance | pneumonia | 2 |
| biocide resistance | child | 1 |
| biocide resistance | kidney | 1 |
| biocide resistance | transplant | 1 |
| biocide resistance | urinary | 2 |
| biocide resistance | MDR | 2 |
| pneumonia | kidney | 1 |
| pneumonia | urinary | 2 |
| child | kidney | 1 |
| child | transplant | 1 |
| child | urinary | 1 |
| child | MDR | 1 |
| kidney | transplant | 1 |
| kidney | urinary | 1 |
| kidney | MDR | 1 |
| transplant | urinary | 1 |
| urinary | MDR | 2 |
| endophytic | root | 2 |
| root | plant | 2 |
| clinical | nosocomial | 1 |
| clinical | bloodstream | 1 |
| clinical | emerging | 1 |
| clinical | growth-promoting | 1 |
| clinical | plant | 1 |
| clinical | water | 1 |
| clinical | root | 1 |
| clinical | larvae | 1 |
| clinical | insect | 1 |
| clinical | gut | 1 |
| clinical | beneficial | 1 |
| clinical | carbapenem-resistant | 1 |
| clinical | endophytic | 1 |
| clinical | animal | 1 |
| clinical | respiratory | 1 |
| clinical | lung | 2 |
| clinical | concern | 1 |
| nosocomial | emerging | 3 |
| nosocomial | growth-promoting | 1 |
| nosocomial | water | 1 |
| nosocomial | root | 1 |
| nosocomial | larvae | 1 |
| nosocomial | gut | 2 |
| nosocomial | carbapenem-resistant | 1 |
| nosocomial | respiratory | 1 |

| | | |
|---|---|---|
| nosocomial | concern | 2 |
| bloodstream | emerging | 1 |
| bloodstream | growth-promoting | 1 |
| bloodstream | water | 1 |
| bloodstream | root | 1 |
| bloodstream | larvae | 1 |
| bloodstream | gut | 1 |
| bloodstream | carbapenem-resistant | 1 |
| bloodstream | respiratory | 1 |
| bloodstream | concern | 1 |
| emerging | growth-promoting | 2 |
| emerging | plant | 2 |
| emerging | water | 1 |
| emerging | root | 1 |
| emerging | larvae | 1 |
| emerging | insect | 2 |
| emerging | gut | 2 |
| emerging | beneficial | 1 |
| emerging | endophytic | 1 |
| emerging | animal | 1 |
| emerging | concern | 2 |
| AMR | growth-promoting | 1 |
| AMR | water | 1 |
| AMR | root | 1 |
| AMR | larvae | 1 |
| AMR | gut | 1 |
| AMR | concern | 2 |
| growth-promoting | water | 1 |
| growth-promoting | root | 1 |
| growth-promoting | larvae | 1 |
| growth-promoting | insect | 1 |
| growth-promoting | gut | 1 |
| growth-promoting | beneficial | 1 |
| growth-promoting | carbapenem-resistant | 1 |
| growth-promoting | animal | 1 |
| growth-promoting | respiratory | 1 |
| growth-promoting | lung | 1 |
| growth-promoting | concern | 1 |
| plant | water | 1 |
| plant | larvae | 1 |
| plant | gut | 1 |
| plant | carbapenem-resistant | 1 |
| plant | respiratory | 1 |
| plant | concern | 1 |
| water | root | 1 |
| water | larvae | 1 |

| | | |
|---|---|---|
| water | insect | 1 |
| water | gut | 2 |
| water | beneficial | 1 |
| water | carbapenem-resistant | 1 |
| water | endophytic | 1 |
| water | animal | 1 |
| water | respiratory | 1 |
| water | lung | 1 |
| water | concern | 1 |
| root | larvae | 1 |
| root | insect | 1 |
| root | gut | 1 |
| root | beneficial | 1 |
| root | carbapenem-resistant | 1 |
| root | animal | 1 |
| root | respiratory | 1 |
| root | lung | 1 |
| root | concern | 1 |
| larvae | insect | 1 |
| larvae | gut | 2 |
| larvae | beneficial | 1 |
| larvae | carbapenem-resistant | 1 |
| larvae | endophytic | 1 |
| larvae | animal | 1 |
| larvae | respiratory | 1 |
| larvae | lung | 1 |
| larvae | concern | 1 |
| insect | gut | 2 |
| insect | carbapenem-resistant | 1 |
| insect | respiratory | 1 |
| insect | concern | 1 |
| gut | beneficial | 1 |
| gut | carbapenem-resistant | 1 |
| gut | endophytic | 1 |
| gut | animal | 1 |
| gut | respiratory | 1 |
| gut | lung | 1 |
| gut | concern | 1 |
| beneficial | carbapenem-resistant | 1 |
| beneficial | respiratory | 1 |
| beneficial | concern | 1 |
| carbapenem-resistant | endophytic | 1 |
| carbapenem-resistant | animal | 1 |
| carbapenem-resistant | respiratory | 1 |
| carbapenem-resistant | lung | 1 |
| carbapenem-resistant | concern | 1 |

| | | |
|---|---|---|
| endophytic | respiratory | 1 |
| endophytic | concern | 1 |
| animal | respiratory | 1 |
| animal | concern | 1 |
| respiratory | concern | 1 |
| lung | concern | 1 |
| nosocomial | environment | 1 |
| nosocomial | opportunistic | 1 |
| nosocomial | animals | 1 |
| nosocomial | healthcare | 2 |
| nosocomial | MDR | 2 |
| nosocomial | biofilm | 2 |
| AMR | animals | 2 |
| AMR | healthcare | 2 |
| AMR | biofilm | 2 |
| emerging | animals | 1 |
| emerging | healthcare | 1 |
| emerging | biofilm | 2 |
| human | animals | 2 |
| human | healthcare | 2 |
| human | concern | 1 |
| human | biofilm | 2 |
| environment | animals | 1 |
| environment | healthcare | 1 |
| environment | concern | 1 |
| environment | biofilm | 1 |
| opportunistic | animals | 1 |
| opportunistic | healthcare | 1 |
| opportunistic | concern | 1 |
| opportunistic | biofilm | 1 |
| animals | healthcare | 1 |
| animals | MDR | 2 |
| animals | concern | 1 |
| animals | biofilm | 1 |
| healthcare | MDR | 2 |
| healthcare | concern | 1 |
| healthcare | biofilm | 2 |
| MDR | concern | 1 |
| MDR | biofilm | 2 |
| concern | biofilm | 1 |
| wound | pneumonia | 1 |
| wound | sepsis | 1 |
| wound | rare | 1 |
| wound | uncommon | 1 |
| wound | compromised | 1 |
| wound | emerging | 1 |

| | | |
|---|---|---|
| wound | NDM-1 | 1 |
| wound | bacteremia | 1 |
| wound | leukemia | 1 |
| wound | insect | 1 |
| wound | bioconversion | 1 |
| wound | industrial | 1 |
| wound | gut | 1 |
| pneumonia | rare | 1 |
| pneumonia | uncommon | 1 |
| pneumonia | compromised | 2 |
| pneumonia | NDM-1 | 1 |
| pneumonia | bioconversion | 1 |
| pneumonia | gut | 1 |
| nosocomial | rare | 1 |
| nosocomial | uncommon | 1 |
| nosocomial | compromised | 2 |
| nosocomial | NDM-1 | 1 |
| nosocomial | bioconversion | 1 |
| nosocomial | industrial | 1 |
| sepsis | rare | 1 |
| sepsis | uncommon | 1 |
| sepsis | compromised | 1 |
| sepsis | emerging | 1 |
| sepsis | NDM-1 | 1 |
| sepsis | bioconversion | 1 |
| sepsis | industrial | 1 |
| sepsis | gut | 1 |
| rare | uncommon | 1 |
| rare | compromised | 1 |
| rare | emerging | 1 |
| rare | NDM-1 | 2 |
| rare | bacteremia | 2 |
| rare | leukemia | 1 |
| rare | insect | 1 |
| rare | bioconversion | 1 |
| rare | industrial | 1 |
| rare | gut | 1 |
| uncommon | compromised | 1 |
| uncommon | emerging | 1 |
| uncommon | NDM-1 | 1 |
| uncommon | bacteremia | 1 |
| uncommon | leukemia | 1 |
| uncommon | insect | 1 |
| uncommon | bioconversion | 1 |
| uncommon | industrial | 1 |
| uncommon | gut | 1 |

| | | |
|---|---|---|
| compromised | emerging | 1 |
| compromised | NDM-1 | 1 |
| compromised | bacteremia | 2 |
| compromised | leukemia | 1 |
| compromised | insect | 1 |
| compromised | bioconversion | 1 |
| compromised | industrial | 1 |
| compromised | gut | 1 |
| emerging | NDM-1 | 1 |
| emerging | bacteremia | 1 |
| emerging | leukemia | 1 |
| emerging | bioconversion | 1 |
| emerging | industrial | 1 |
| NDM-1 | bacteremia | 2 |
| NDM-1 | leukemia | 1 |
| NDM-1 | insect | 1 |
| NDM-1 | bioconversion | 1 |
| NDM-1 | industrial | 1 |
| NDM-1 | gut | 1 |
| bacteremia | bioconversion | 1 |
| bacteremia | industrial | 1 |
| bacteremia | gut | 1 |
| leukemia | bioconversion | 1 |
| leukemia | industrial | 1 |
| leukemia | gut | 1 |
| insect | bioconversion | 1 |
| insect | industrial | 1 |
| bioconversion | industrial | 1 |
| bioconversion | gut | 1 |
| industrial | gut | 1 |
| crops | emerging | 2 |
| crops | plant | 1 |
| crops | human | 1 |
| crops | growth-promoting | 1 |
| crops | potato | 2 |
| crops | biocontrol | 1 |
| crops | antifungal | 1 |
| crops | rice | 2 |
| crops | leaves | 1 |
| crops | banana | 2 |
| crops | rot | 2 |
| emerging | potato | 2 |
| emerging | biocontrol | 1 |
| emerging | antifungal | 1 |
| emerging | rice | 2 |
| emerging | leaves | 1 |

| | | |
|---|---|---|
| emerging | banana | 2 |
| emerging | rot | 2 |
| plant | potato | 1 |
| plant | rice | 1 |
| plant | rot | 1 |
| human | growth-promoting | 1 |
| human | potato | 1 |
| human | rice | 1 |
| human | banana | 1 |
| human | rot | 1 |
| growth-promoting | potato | 1 |
| growth-promoting | biocontrol | 1 |
| growth-promoting | antifungal | 1 |
| growth-promoting | rice | 1 |
| growth-promoting | leaves | 1 |
| growth-promoting | banana | 1 |
| growth-promoting | rot | 1 |
| potato | biocontrol | 1 |
| potato | antifungal | 1 |
| potato | rice | 2 |
| potato | leaves | 1 |
| potato | banana | 2 |
| potato | rot | 2 |
| biocontrol | rice | 1 |
| biocontrol | banana | 1 |
| biocontrol | rot | 1 |
| antifungal | rice | 1 |
| antifungal | banana | 1 |
| antifungal | rot | 1 |
| rice | leaves | 1 |
| rice | banana | 2 |
| rice | rot | 2 |
| leaves | banana | 1 |
| leaves | rot | 1 |
| banana | rot | 2 |
| AMR | rare | 1 |
| AMR | carbapenemase | 1 |
| AMR | NDM-1 | 1 |
| human | rare | 1 |
| human | carbapenemase | 1 |
| human | NDM-1 | 1 |
| MDR | rare | 1 |
| MDR | bacteremia | 2 |
| MDR | carbapenemase | 1 |
| MDR | NDM-1 | 1 |
| MDR | blood | 1 |

| | | |
|---|---|---|
| animals | rare | 1 |
| animals | clinical | 1 |
| animals | bacteremia | 1 |
| animals | carbapenemase | 1 |
| animals | NDM-1 | 1 |
| animals | lung | 1 |
| animals | blood | 1 |
| rare | clinical | 1 |
| rare | carbapenemase | 1 |
| rare | lung | 1 |
| rare | blood | 1 |
| clinical | bacteremia | 1 |
| clinical | carbapenemase | 1 |
| clinical | NDM-1 | 1 |
| clinical | blood | 1 |
| bacteremia | carbapenemase | 1 |
| carbapenemase | NDM-1 | 1 |
| carbapenemase | lung | 1 |
| carbapenemase | blood | 1 |
| NDM-1 | lung | 1 |
| NDM-1 | blood | 1 |
| crops | biofilm | 1 |
| rice | biofilm | 1 |
| rot | biofilm | 1 |
| banana | biofilm | 1 |
| biofilm | potato | 1 |
| antifungal | rhizosphere | 1 |
| human | rhizosphere | 1 |
| human | biocide resistance | 1 |
| human | compromised | 1 |
| human | urinary | 1 |
| human | threat | 1 |
| cystic fibrosis | biofilm | 1 |
| cystic fibrosis | rhizosphere | 1 |
| cystic fibrosis | biocide resistance | 1 |
| cystic fibrosis | compromised | 1 |
| cystic fibrosis | urinary | 1 |
| cystic fibrosis | threat | 1 |
| cystic fibrosis | healthcare | 1 |
| AMR | rhizosphere | 1 |
| AMR | compromised | 1 |
| biofilm | rhizosphere | 1 |
| biofilm | animal | 1 |
| biofilm | biocide resistance | 1 |
| biofilm | compromised | 1 |
| biofilm | pneumonia | 1 |

| | | |
|---|---|---|
| biofilm | urinary | 1 |
| biofilm | bacteremia | 1 |
| biofilm | chronic | 1 |
| biofilm | lung | 1 |
| biofilm | threat | 1 |
| biofilm | immunocompromised | 1 |
| rhizosphere | animal | 1 |
| rhizosphere | biocide resistance | 1 |
| rhizosphere | compromised | 1 |
| rhizosphere | nosocomial | 1 |
| rhizosphere | pneumonia | 1 |
| rhizosphere | urinary | 1 |
| rhizosphere | bacteremia | 1 |
| rhizosphere | chronic | 1 |
| rhizosphere | MDR | 1 |
| rhizosphere | lung | 1 |
| rhizosphere | threat | 1 |
| rhizosphere | healthcare | 1 |
| rhizosphere | immunocompromised | 1 |
| animal | biocide resistance | 1 |
| animal | compromised | 1 |
| animal | urinary | 1 |
| animal | chronic | 1 |
| animal | MDR | 1 |
| animal | threat | 1 |
| animal | healthcare | 1 |
| biocide resistance | compromised | 1 |
| biocide resistance | nosocomial | 1 |
| biocide resistance | bacteremia | 1 |
| biocide resistance | chronic | 1 |
| biocide resistance | lung | 1 |
| biocide resistance | threat | 1 |
| biocide resistance | healthcare | 1 |
| biocide resistance | immunocompromised | 1 |
| compromised | urinary | 1 |
| compromised | chronic | 1 |
| compromised | MDR | 1 |
| compromised | lung | 1 |
| compromised | threat | 1 |
| compromised | healthcare | 1 |
| compromised | immunocompromised | 1 |
| nosocomial | urinary | 1 |
| nosocomial | chronic | 1 |
| nosocomial | threat | 1 |
| pneumonia | chronic | 1 |
| pneumonia | threat | 1 |

| | | |
|---|---|---|
| pneumonia | healthcare | 1 |
| urinary | bacteremia | 1 |
| urinary | chronic | 1 |
| urinary | lung | 1 |
| urinary | threat | 1 |
| urinary | healthcare | 1 |
| urinary | immunocompromised | 1 |
| bacteremia | chronic | 1 |
| bacteremia | threat | 1 |
| bacteremia | healthcare | 1 |
| chronic | healthcare | 1 |
| chronic | immunocompromised | 1 |
| MDR | threat | 1 |
| MDR | immunocompromised | 1 |
| lung | threat | 1 |
| lung | healthcare | 1 |
| threat | healthcare | 1 |
| threat | immunocompromised | 1 |
| healthcare | immunocompromised | 1 |
| root | soil | 1 |
| root | antimicrobial | 1 |
| root | antifungal | 1 |
| soil | antifungal | 1 |
| antimicrobial | antifungal | 1 |
| aquaculture | water | 1 |
| gut | aquaculture | 1 |
| gut | cultured | 2 |
| gut | epizootic | 1 |
| cultured | water | 1 |
| epizootic | water | 1 |
| larvae | healthy | 1 |
| larvae | cultured | 1 |
| gut | healthy | 2 |
| healthy | cultured | 2 |
| healthy | fish | 1 |

**Table S4.** Table of bacterial LOXs included in the binding site bottom statistical analysis. The table provides the species which carry corresponding LOXs, binding site bottom residues (aligned to the corresponding residues in *Pseudomonas aeruginosa* in the header row), and the volume sums of these residues
(given in $10^{-3}$ $nm^3$)

| **Species/strain** | **Glu369** | **Met434** | **Ile608** | **Volume sum** |
|---|---|---|---|---|
| *Pantoea sp. OXWO6B1* | K | L | I | 506,8 |
| *Kosakonia sp. AG348* | K | L | I | 506,8 |

| | | | | |
|---|---|---|---|---|
| *Pluralibacter gergoviae* | K | L | I | 506,8 |
| *Nocardia brasiliensis* | F | I | V | 494,6 |
| *Burkholderia singularis* | E | L | I | 475 |
| *Moellerella wisconsensis* | E | L | I | 475 |
| *Variovorax paradoxus* | F | I | L | 523,8 |
| *Mycobacteroides abscessus* | N | L | T | 406,4 |
| *Nocardia pseudobrasiliensis* | F | I | V | 494,6 |
| *Pseudomonas aeruginosa* | E | M | I | 472,2 |
| *Kutzneria sp. 744* | N | L | V | 426,9 |
| *Variovorax gossypii* | F | I | L | 523,8 |
| *Burkholderia thailandensis* | E | L | I | 475 |
| *Variovorax guangxiensis* | F | I | L | 523,8 |
| *Nocardia seriolae* | F | I | V | 494,6 |
| *Burkholderia gladioli* | E | L | I | 475 |
| *Kosakonia sacchari* | K | L | I | 506,8 |
| *Lentzea kentuckyensis* | N | L | V | 426,9 |
| *Pseudobacteriovorax antillogorgiicola* | S | L | T | 378 |
| *Pantoea ananas* | K | L | I | 506,8 |
| *Colwellia echini* | S | D | F | 396,8 |
| *Enterovibrio norvegicus* | C | V | F | 433,9 |
| *Shewanella waksmanii* | C | V | F | 433,9 |
| *Algibacillus agarilyticus* | C | N | F | 415,2 |
| *Aliikangiella coralliicola* | C | T | F | 413,4 |
