## Supplementary Figures for "Bacterial lipoxygenases are associated with host-microbe interactions and may provide cross-kingdom host jumps"

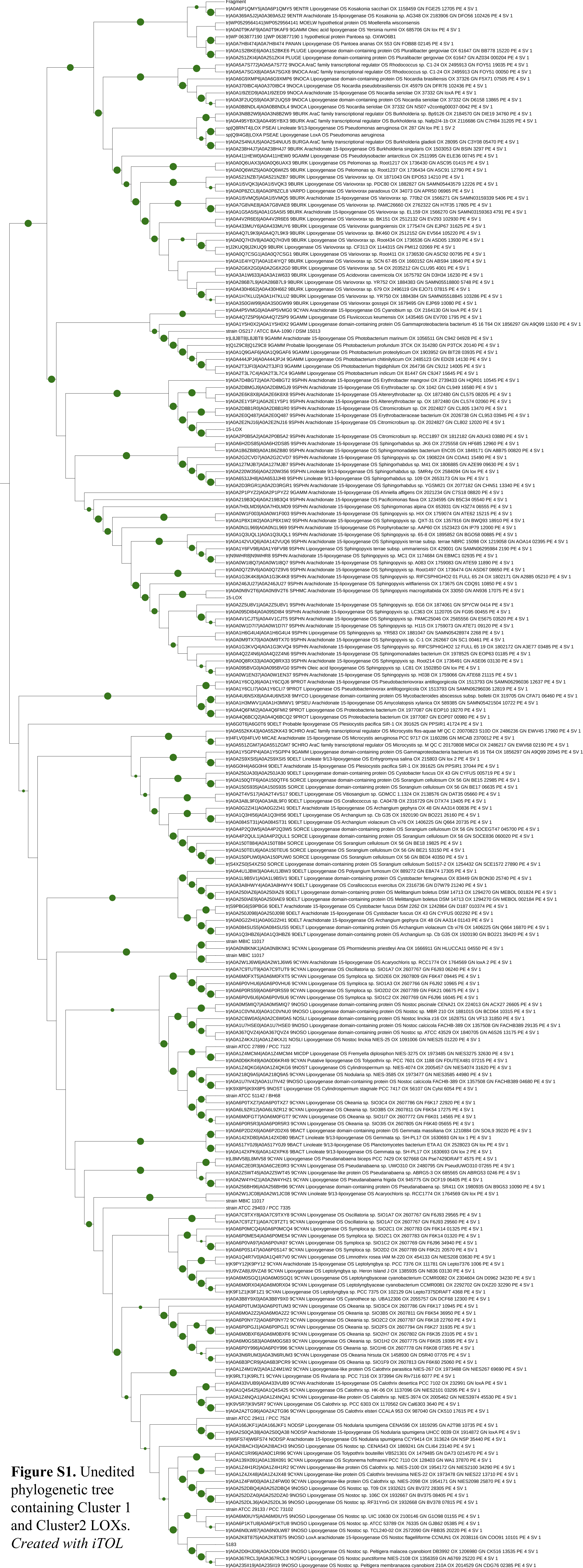

**Figure S1.** Unedited phylogenetic tree containing Cluster 1 and Cluster2 LOXs. *Created with iTOL*

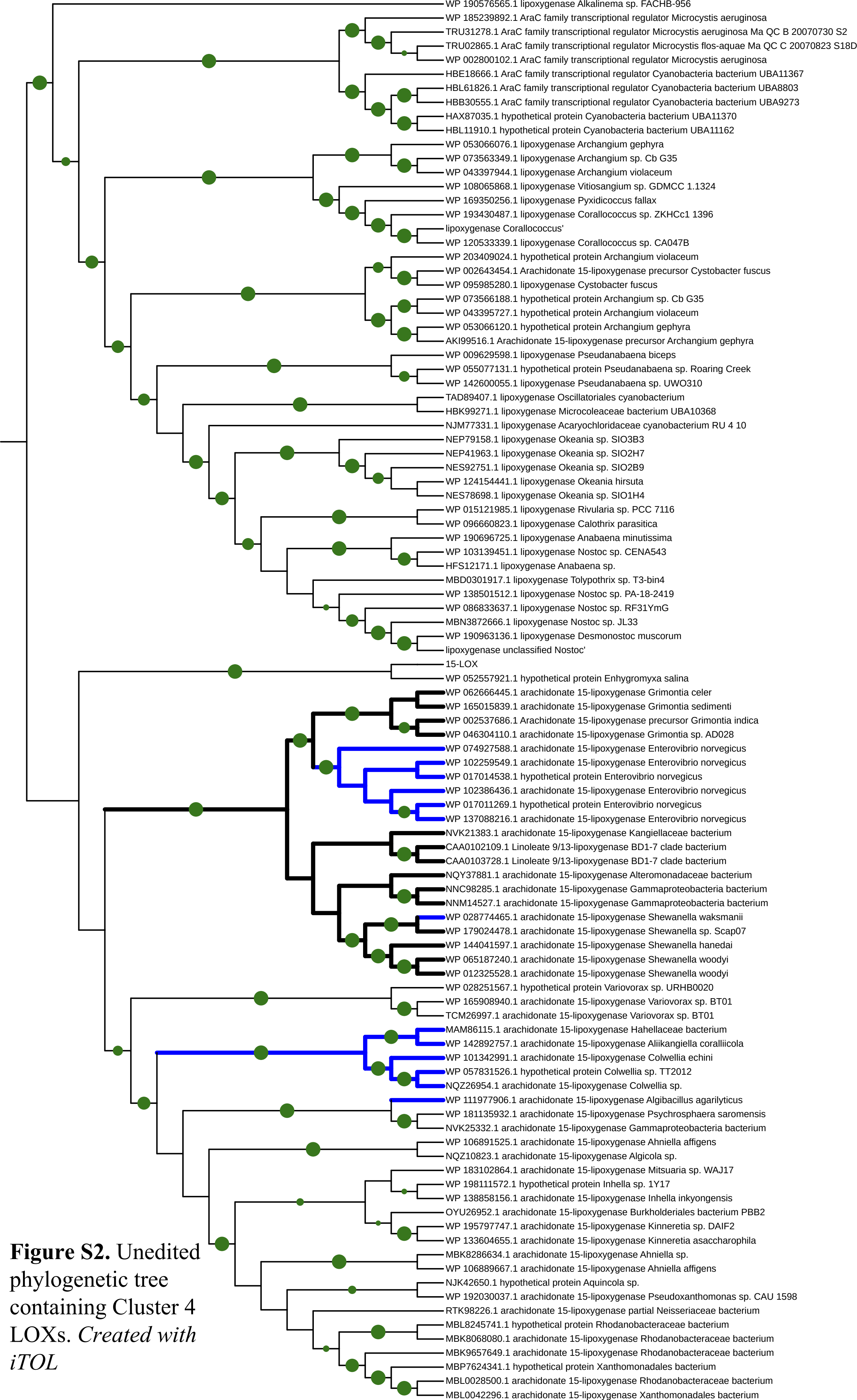


**Figure S2.** Unedited phylogenetic tree containing Cluster 4 LOXs. *Created with iTOL*

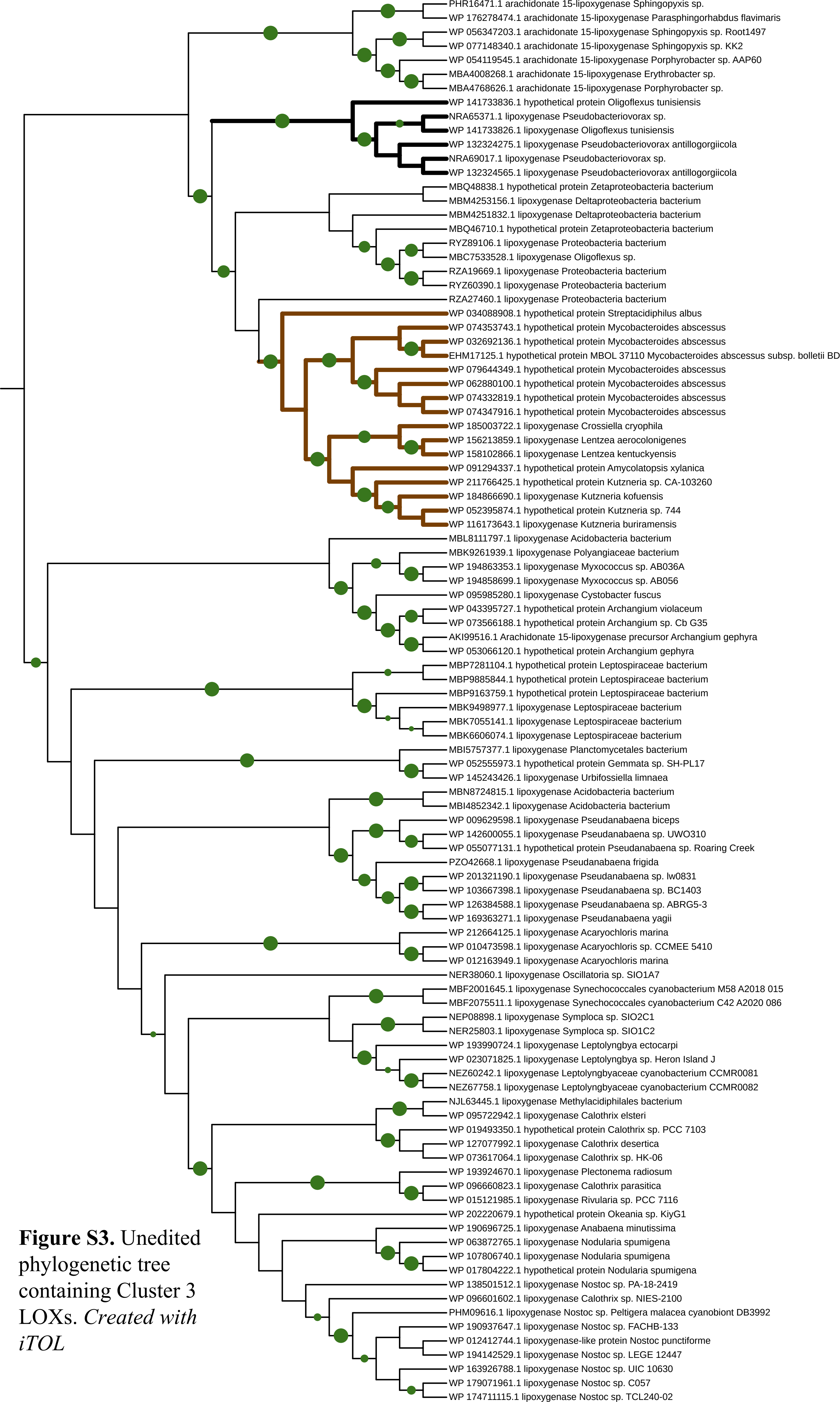


**Figure S3.** Unedited phylogenetic tree containing Cluster 3 LOXs. *Created with iTOL*
